# Induction of ferroptotic and amyloidogenic signatures linked to Alzheimer’s disease by chemically distinct air pollutants

**DOI:** 10.64898/2026.01.06.696601

**Authors:** Kristina Shkirkova, Naomi S. Sta Maria, Herbert Anson, Yashar Aghaei, Mohammad Mahdi Badami, Ararat Chakhoyan, Jose A. Godoy-Lugo, Claire Chung, Salma Durra, Angela Tang-Tan, Lifu Zhao, Alexandra Demetriou, Manuel Morales, Sindhu Daggupati, Isabella Bent, Hyoungjin Park, Caleb Franklin, Selena Chen, Giovanni Chahine, Skyye Dodds-Lewis, Masako Morishita, Russell E. Jacobs, Jean-François Gout, Wendy J. Mack, Henry Jay Forman, Bérénice A. Benayoun, Marc Vermulst, Mitchell D. Cohen, Matthew Campen, Constantinos Sioutas, William J. Mack, Caleb E. Finch, Max A. Thorwald

## Abstract

Air pollution (AirP) exposure is associated with increased Alzheimer’s disease (AD) risk, yet AirP is chemically heterogeneous, complicating identification of shared pathogenic drivers. We examined acute cortical responses to two metal-rich AirP sources, diesel exhaust particles (DEP) and World Trade Center (WTC) dust, and compared them to woodsmoke (WS), a particulate exposure with low metal content. DEP and WTC elicited highly convergent transcriptional responses, sharing over 1200 differentially expressed genes linked to inflammation, ferroptosis, neuronal remodeling, and amyloid processing. These changes were accompanied by impaired antioxidant activity and increased lipid peroxidation within lipid rafts, a membrane microdomain critical for amyloid processing, resulting in increased Aβ generation. In contrast, WS produced a distinct transcriptional signature and failed to induce ferroptotic priming or lipid peroxidation, consistent with its low metal composition. Together, these findings implicate metals as a shared driver linking diverse AirP exposures to amyloidogenic vulnerability and elevated AD risk.

**Graphical Abstract:** Acute AirP exposure converges on ferroptotic priming, amyloidogenic processing, and white-matter vulnerability. Acute exposure to metal-rich AirP, such as DEP or WTC introduces redox-active metals and particulate matter that promote lipid peroxidation, amyloidogenesis, and altered transcriptional regulation in the brain. AirP exposure engages xenobiotic metabolism pathways (AhR/ARNT), activates iron and heme handling through ferritinophagy (NCOA4) and heme oxygenase activity (HMOX1), and blunts lipid peroxide detoxification systems, including glutathione peroxidase 4 (GPx4), ferroptosis suppressor protein 1 (FSP1), and glutathione (GSH) synthesis. These changes promote ferroptotic priming and lipid raft oxidation, facilitating amyloid precursor protein (APP) processing by secretases (ADAM10, BACE1, γ-secretase) and increasing amyloid-β (Aβ) generation. In parallel, transcriptional and cell-state remodeling involving neuronal and oligodendrocyte responses contribute to selective white-matter vulnerability, particularly within the corpus callosum. Together, these pathways provide a mechanistic framework linking acute AirP exposure to convergent oxidative, amyloidogenic, and microstructural changes relevant to Alzheimer’s disease pathology.

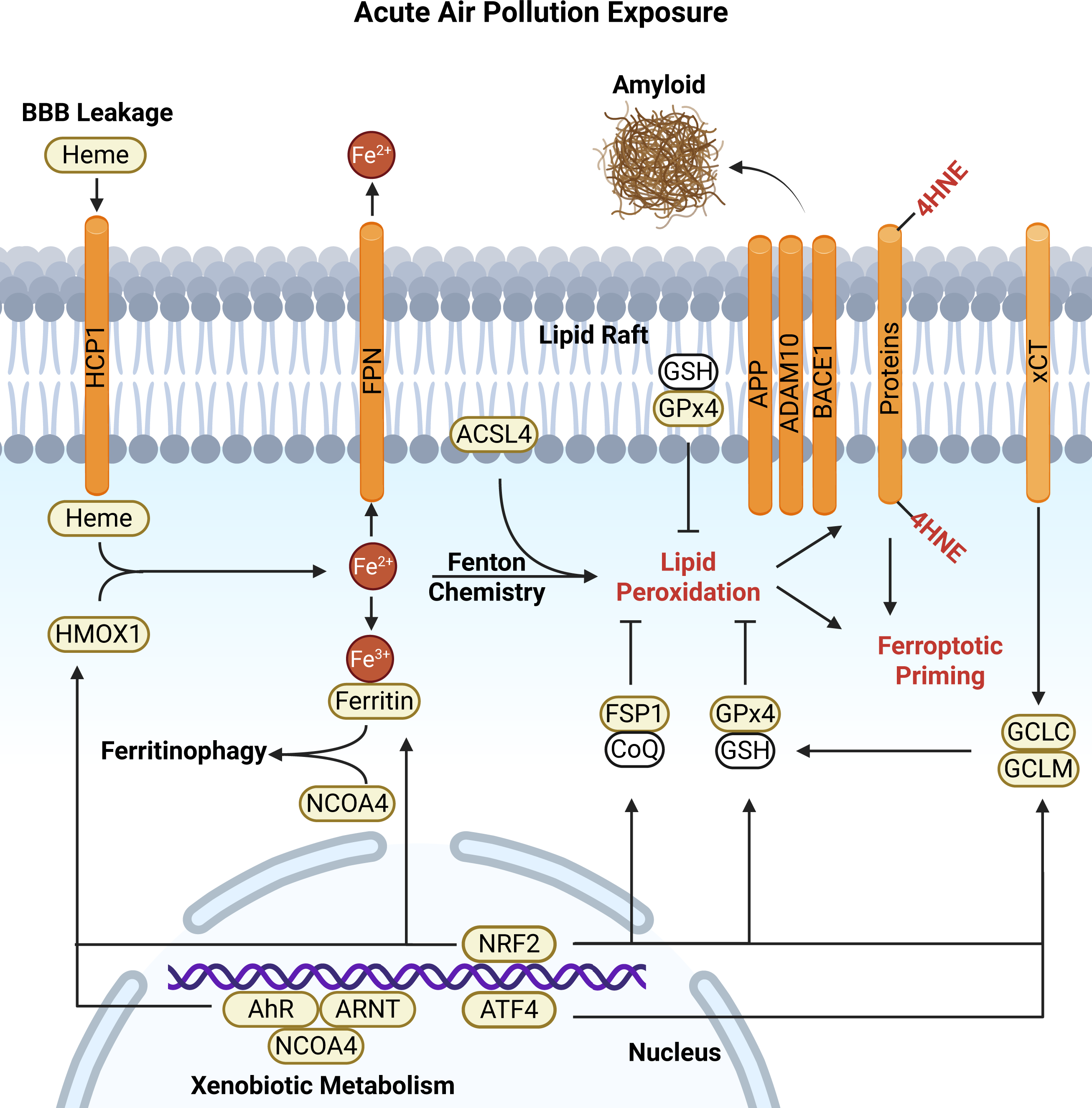

## INTRODUCTION

AD is a progressive neurodegenerative disorder characterized by memory loss and cognitive decline. While 30 million people worldwide are currently affected, the increase of life expectancy projects a fivefold increase of AD by 2050^1^. Although the molecular mechanisms underlying AD are incompletely understood, aberrant processing and aggregation of β-amyloid (Aβ) peptides are widely implicated in AD progression^2^. Aβ oligomers, fibrils, and plaques disrupt synaptic function, promote oxidative injury, and contribute to neuronal dysfunction and loss^3^. While genetic risk factors and aging are well-established contributors to AD, growing evidence indicates that environmental exposures also play a significant role in modulating disease risk and progression.

Among environmental factors, air pollution (AirP) has emerged as a prominent and independent risk factor for dementia and AD^4–7^. Epidemiological studies across diverse populations consistently demonstrate that chronic exposure to particulate matter (PM) and traffic-related air pollution (TRAP) is associated with increased dementia incidence, accelerated cognitive decline, and earlier AD onset^8–11^. Importantly, these associations persist after adjustment for socioeconomic status, education, and lifestyle factors, underscoring AirP as a modifiable contributor to neurodegenerative disease. Despite this strong epidemiological signal, the biological mechanisms linking AirP exposure to AD pathology remain poorly defined.

A major challenge in understanding AirP-associated neurodegeneration lies in the chemical heterogeneity of AirP itself^12^. AirP encompasses a complex mixture of PM and gases originating from diverse sources, including traffic emissions, industrial activity, structural fires, and biomass combustion. Consequently, it remains unclear whether neurodegenerative outcomes associated with AirP arise from a shared biological response or from exposure-specific mechanisms. Notably, many AirP sources contain substantial quantities of metals, including Fe, Cu, Zn, Al, Pb, Cd, Hg, and Mn, which are independently associated with neurotoxicity and oxidative stress^13–15^. Unlike reactive gaseous pollutants such as ozone or NO_2_, which primarily exert transient effects in peripheral tissues^16^, metals can accumulate within the brain parenchyma, persist for extended periods, and continuously redox-cycle, sustaining oxidative burden^17–19^.

Brain metal accumulation can occur through multiple routes, including blood-brain barrier (BBB) dysfunction, active transport mechanisms, and cerebral microhemorrhages (CMBs)^20,21^. CMBs arise decades before clinical AD, with increasing prevalence beginning in early adulthood, and are markedly elevated in AD brains^22,23^. These microvascular events permit the extravasation of heme-bound iron into the brain parenchyma, where iron accumulates within plaques and surrounding tissue. Because CMBs frequently localize to periventricular and subcortical white-matter regions, iron deposition arising from microvascular injury is positioned to disproportionately impact white-matter tracts, including the corpus callosum, which exhibits early vulnerability in AirP-associated neurotoxicity^24^. Iron-laden amyloid plaques are a consistent feature of AD neuropathology and provide a catalytic environment for lipid peroxidation and oxidative damage^25–27^. Together, these observations position persistent brain metal burden as a plausible convergent driver linking chemically distinct AirP exposures to AD-relevant pathology.

We and others have highlighted iron-dependent lipid peroxidation and ferroptosis-associated pathways as important contributors to neurodegeneration, including AD^18,26,27^. Iron readily cycles between ferrous (Fe^2+^) and ferric (Fe^3+^) states, generating highly reactive hydroxyl radicals capable of indiscriminate molecular damage^28^. Because membrane lipids represent a major target of iron-catalyzed oxidation, sustained lipid peroxidation can overwhelm antioxidant defenses and compromise membrane integrity. When lipid peroxide detoxification systems, most notably glutathione peroxidase 4 (GPx4), are impaired, or glutathione (GSH) availability is limited, cells enter a ferroptosis-destined state^29^. While definitive demonstration of ferroptotic cell death in human AD remains challenging, multiple convergent biochemical and molecular markers strongly support ferroptosis-associated processes as a component of AD pathology^18,26,27,30–32^.

Within neuronal membranes, lipid rafts (LRs) represent microdomains enriched in polyunsaturated lipids and cholesterol that are particularly vulnerable to oxidative damage. Importantly, LRs also harbor the secretase complexes responsible for amyloid precursor protein (APP) processing, positioning them as a potential mechanistic nexus between iron-mediated lipid peroxidation and amyloidogenic processing. Analyses of postmortem human AD brain tissue demonstrate diminished GPx4 and GPx1 protein levels and activity within cortical LRs, accompanied by increased lipid peroxidation^26,27^. Decomposition of oxidized lipids generates reactive aldehydes such as 4-hydroxynonenal (HNE), which form irreversible protein adducts that alter enzymatic function^33,34^. HNE modifications have been shown to favor pro-amyloidogenic APP processing, directly linking lipid oxidation within LRs to increased Aβ production^35,36^.

Despite these insights, it remains unclear whether chemically distinct AirP sources converge on shared molecular pathways relevant to AD, or whether neurodegenerative risk reflects exposure-specific mechanisms. To address this gap, we directly compared three well described environmentally and chemically distinct AirP exposures: diesel exhaust particles (DEP)^12,24,37,38^, a canonical component of TRAP; dust generated by the World Trade Center (WTC) terrorist attacks, which has been associated with an approximately decade-earlier AD trajectory^39–42^; and woodsmoke (WS), a biomass-derived pollutant of increasing relevance due to the growing frequency and severity of wildfires, including recent events in the Los Angeles region^43–45^. By examining whether these exposures converge or diverge in their ability to promote AD-associated molecular, biochemical, and microstructural changes, this study tests the hypothesis that persistent metal-mediated oxidative stress represents a unifying mechanism linking AirP exposure to AD pathology.

Here, we show that acute exposure to chemically distinct, metal-rich AirP sources, DEP and WTC, elicits convergent molecular responses characterized by oxidative stress, impaired lipid peroxide detoxification, ferroptosis-associated priming, altered amyloidogenic processing, and preferential vulnerability of white-matter tracts. Despite marked differences in chemical composition, both exposures engage shared transcriptional, biochemical, and microstructural pathways relevant to AD pathology. In contrast, WS exposure induces a divergent transcriptional and biochemical profile that does not recapitulate these convergent features. Together, these findings identify metal-associated oxidative mechanisms as a common biological link between specific forms of AirP and AD relevant pathology and distinguishes them from other particulate exposures.

## METHODS

### Chemical Composition analysis

DEP and WTC chemical composition analysis was performed at the Desert Research Institute’s environmental analysis facility. The measurement of OC and EC was conducted using a multiwavelength thermal/optical carbon analyzer (Magee Scientific, Berkeley, CA, USA), adhering to the protocols delineated by the interagency monitoring of protected visual environments^46^. Inorganic ion content was conducted by ion chromatography (IC) by eluting the particles into ultrapure deionized water using sonication, followed by filtration and analysis as previously described ^47,48^. Inductively coupled plasma mass spectroscopy (ICP-MS) was employed to determine the concentrations of metals and trace elements, involving a hot block acid digestion process to extract by nitric (HNO_3_), hydrochloric (HCl), and hydrofluoric (HF) acid ^49,50^. After digestion, the samples were diluted with deionized water, aerosolized, and then introduced into the ICP-MS instrument (Thermo Finnigan Element 2, Thermo Fisher Scientific Inc., Waltham, MA, USA). Woodsmoke metal analysis was also performed by ICP-MS at Michigan State University. Chemical composition and particle sizes for DEP, WTC, and WS have been previously described^41,43,51^.

### Animal

Procedures were reviewed and approved by the University of Southern California Institutional Animal Care and Use Committees (IACUC) under protocol #20842. Male and female mice C57BL/6 (n=10/group/sex) were housed separately in groups of five, at 22°C/30% humidity and 12 hours of light-dark cycles with standard nesting, food, and water ad libitum. 2-month-old mice were exposed to re-aerosolized DEP (NIST SRM 2975), WTC dust, or control filtered air (FA) for 5 hours at 100 μg/m^3^. Briefly, DEP or WTC dust were Milli-Q deionized water-solubilized, mixed, and sonicated for 30 minutes for re-aerosolized animal exposure^37^. After the 5-hour exposure, mice were humanly euthanized and perfused with PBS. One hemisphere of the brain was micro-dissected, frozen, and then stored at -80°C, second hemisphere was fixed in 4% PFA for 24 hours and embedded in paraffin for immunofluorescent analysis. Four whole brains per sex in each exposure group (n=4) were saved for *ex vivo* MRI.

For the WS exposure, animal procedures were approved by the University of New Mexico IACUC. Female C57BL/6J mice at 2 months of age were exposed to laboratory-generated WS (500µg/m^3^) for 4 hours. WS was derived from cedar chips harvested in the Blue Gap Tachee, Arizona region^44^. Post-exposure, mice were euthanized and perfused with PBS. One hemisphere was micro-dissected and stored at -80°C, the second hemisphere was frozen in -80°C and embedded in OCT for immunofluorescent analysis on frozen sections.

### MRI acquisition

Was performed at the Biological Imaging and Spectroscopy Core at the Zilkha Neurogenetic Institute (University of Southern California). The cryogen-free MR Solutions PET/MRI 7T system was equipped with a bore size of ∼24 cm, a maximum gradient up to 600 mT/m, and a 20 mm internal diameter quadrature bird cage mouse head coil. *Ex vivo* fixed brains with the skulls intact were prepared following cardiac perfusion and were incubated in 5 mM Gd-DTPA (BioPal, Inc.) for 3 days. The samples were then immersed in perfluorinated polyether fluid (Galden) and positioned with a rubber silicone spacer in a 15 mL centrifuge tube prior to MRI.

A positioning gradient echo sequence was first acquired to prepare the slice stacks for the 3D multi-echo multi-slice (MEMS) spin echo sequence. The MEMS parameters are as follows: echo times (TEs) = 10, 20, 30, 40, 50, 60, 70, 80, 90, and 100 ms; TR = 176 ms; slice thickness = 0.6 mm; field of view (FOV) = 18 mm x 18 mm x 19.2 mm; acquired matrix size (MS) = 256 x 128 x 64 and resized to 192 x 192 x 32; flip angle (FA) = 50°; acquisition bandwidth = 50 kHz; and NA = 1.

2D spin echo diffusion weighted imaging (SEDWI) was performed calculate diffusion and fractional anisotropy maps and included the following parameters: TE = 23 ms, TR = 4000 ms, 3 b-values (600, 1200, and 1800 s/mm^2^), 20 directions, FOV = 18 mm x 18 mm, slice thickness = 0.6 mm, 32 slices, MS = 150 x 150, and acquisition bandwidth = 50kHz.

### MRI Analysis

T_2_ maps were generated from the MEMS images through a pixel-by-pixel exponential fitting of the signal intensities across the different TE times. MATLAB Rocketship v.1.4 module was used to perform the parametric fits^52^. All fits with an r^2^>0.6 were included. Using Fiji software, pixels with fits having r^2^ values <0.6 were set to not-a-number (NaN, i.e., missing data) and were not included in the analysis. Additionally, brain regions were extracted by manually delineating brain outlines on each slice, and outside brain regions were set to NaN. R_2_ maps were generated from the T_2_ maps, using the relationship T_2_ = 1/R_2_. Mean R_2_ values for each ROI, for each subject, were obtained. Apparent diffusion coefficient (ADC) and fractional anisotropy (FA) maps were generated from the SEDWI scans using DSI studio software^53^. Regions of interest (ROI) were manually delineated using the polygon tool in Fiji.

### Western Blot

Lipid rafts, nuclear lysates, and whole cell lysates (RIPA) were generated from mouse cortex as previously described and validated^26,27,38,54,55^. Whole cell lysates (20μg), lipid rafts (5μg), or nuclear lysates (20ug) were boiled at 75°C under denatured conditions and resolved on 4-20% gradient gels. Proteins were electroblotted using a Criterion blotter (Bio-Rad Laboratories, Hercules, CA) and transferred onto 0.45μm polyvinyl difluoride membranes. Membranes were stained using Revert 700 fluorescent protein stain and imaged before blocking with LI-COR Intercept blocking buffer (LI-COR Biosciences, Lincoln, NE), followed by primary antibody incubation overnight at 4°C. Membranes incubated with IRDye 800CW and/or 700CW secondary antibodies and visualized by Odyssey (LI-COR Biosciences). Western blot data were quantified with ImageJ and normalized by total protein per lane and/or loading control protein.

### Dot blot

RIPA (20μg) or lipid raft (5μg) lysates were loaded onto a dot blot apparatus for gravity filtration for 3 hours. After filtration, membranes were strained with Revert 700 fluorescent protein stain and imaged before blocking with LI-COR Intercept blocking buffer (LI-COR Biosciences, Lincoln, NE), followed by primary antibody incubation overnight at 4°C.

### Immunofluorescence

For DEP and WTC exposures, paraffin-embedded brains were sectioned at 5µm thickness and mounted on slides. Sections were deparaffinized and rehydrated, followed by antigen retrieval using sodium citrate buffer. Slides were then blocked with blocking solution and incubated with primary antibodies overnight at 4 °C, followed by incubation with appropriate secondary antibodies for 1 hour the following day. Paraffin-embedded sections encompassing the corpus callosum and cortex were stained for markers of lipid peroxidation (HNE) and DNA oxidation (8-OHdG).

For WS, frozen OCT-embedded brains were sectioned at 15µm thickness and mounted on slides. Slides were blocked with donkey serum and incubated with primary and secondary antibodies. Frozen brain sections encompassing the corpus callosum and hippocampus were stained for markers of degraded myelin (dMBP), oxidative damage (HNE, 8-OHdG), complement activation (C5, C5a), and microglial reactivity (Iba1).

To quantify immunofluorescent signal in both frozen and paraffin-embedded tissue, brain sections were imaged at 40× magnification using a BZ-X810 Keyence microscope (Keyence, USA). Image analysis for each brain region was performed by two observers blinded to exposure group using NIH ImageJ software. Representative images were selected for display and adjusted uniformly for brightness and contrast across all groups.

### Biochemical Assays

Total protein was quantified by 660nm assay for lipid rafts and nuclear lysate, and BCA for whole cell lysates (Thermo Fisher Scientific, Waltham, MA). Total glutathione peroxidase activity was calculated by activity assay (Cayman Chemical, Ann Arbor, MI). For phospholipid hydroperoxidase activity, oxidized phosphatidyl choline (PCOOH) was generated^56^ and used in place of the provided cumene hydroperoxide^26,27,57^. The same batch of PCOOH was utilized for all assays. Tissue heme levels were quantified by Quantichrom assay (BioAssay Systems, Hayward, CA). All optical densities were measured using a SpectraMax M2 spectrophotometer equipped with a temperature regulator (Molecular Devices, San Jose, CA).

### RNA & Circle sequencing

20 mg of brain tissue was homogenized in TRIzol reagent using a BeadBug Benchtop Homogenizer. Brain tissue was suspended in 1 mL of TRIzol and homogenized for 6 rounds of 10 seconds homogenization, and 60-second holds between each round. Following homogenization, 300 µL of chloroform was added, and samples were centrifuged with heavy gel phase-lock tubes (VWR, 10847-802) to separate the aqueous phase. The aqueous phase was applied to a standard column-based RNA purification kit (Quantabio, Extracta Plus, Cat# 95214-050) following the manufacturer’s protocol. RNA concentration and integrity were assessed with a Qubit 4.0 Fluorometer and Agilent Bioanalyzer prior to sequencing. Library preparation (mRNA library, poly A enrichment) and RNA sequencing (NovoSeq PE150. 6G raw data per sample) was performed by Novogene. Reads were trimmed with trim_galore-0.6.5-1 and mapped to the Mus musculus GRCm39 reference genome with STAR-2.7.3a^58^. Mapped reads were counted to genes using featureCounts (Subread-2.0.3)^59^. Differential expression analysis was performed with DESeq2-1.34.0^60^. For DEP/WTC exposure datasets, sex was included as a covariate. For the WS dataset, expression data exhibited dominant variance unrelated to exposure status, as assessed by principal component analysis. To improve detection of exposure-associated transcriptional changes, residual technical and latent biological variation was adjusted using RUVSeq (v1.32.0). RUV correction was not applied to DEP or WTC datasets, which showed strong exposure-driven separation without adjustment. Statistical significance for differential expression was defined using Benjamini-Hochberg false discovery rate-adjusted p-values (FDR<0.05). DEGs were subjected to Gene set enrichment analysis (GSEA) analysis for pathway enrichment. Circle-sequencing was performed as previously described with our error detection pipeline^61,62^.

To determine the influence of cellular composition effects, estimated cell-type proportions were derived from bulk RNA-seq using MuSiC v1.1.0^63^ using single-cell reference datasets from Celldex; MouseBrainData, and MouseRNAseqData. Estimated cell proportions were incorporated as covariates in reanalysis by DESeq2 to isolate transcriptomic effects independent of compositional differences among exposure groups. Transcription factor activity was inferred using DoRothEA V1.6.9 by high-confidence regulon and VIPER v1.28.0 for enrichment scoring through full ranked gene lists as inputs. Visualization of RNA data by upset, PCA, or hexbin plots used UpSetR, ggplot2, pheatmap, and RColorBrewer. Raw RNA-seq data are available through Annotare: E-MTAB-16199.

### Statistics

Were performed using SPSS (Version 29.0.20.0; IBM, Chicago, IL). For the majority of biochemical, molecular, and transcriptomic endpoints, means among exposure groups were compared using analysis of covariance (ANCOVA) with sex included as a covariate to adjust for sex-related differences among study groups. Pairwise comparisons were performed using Bonferroni correction with significance set at p<0.05. Homogeneity of variances was assessed using Levene’s test, and non-normally distributed data were log-transformed prior to analysis when appropriate. MEMS and DTI measures were analyzed using two-way ANOVA to assess the effects of exposure and brain region, followed by Tukey’s post hoc testing for multiple comparisons. Because regional differences were not the biological focus, post hoc testing and reporting were restricted to within-region exposure comparisons; cross-region comparisons were not performed or reported. For WS analyses, comparisons were performed between WS exposed and FA groups using two-tailed unpaired t-tests. Data are presented as mean ± SEM unless otherwise indicated.

## RESULTS

### Distinct chemical signatures of DEP and WTC converge on shared metal enrichment

Comprehensive chemical profiling by ion chromatography, thermal-optical analysis, and inductively coupled plasma mass spectrometry (ICP-MS) revealed that DEP and WTC dust exhibit fundamentally distinct ionic, carbonaceous, and size characteristics, but converge on enrichment of redox-active transition metals. DEP was enriched in NH_4_^+^ and SO ^2-^ and contained high elemental and organic carbon content, consistent with combustion-derived exhaust, whereas WTC dust showed elevated levels of soluble Cl^-^, Na^+^, and K^+^, consistent with pulverized building materials (**S.Fig. 1A,B**).

Furthermore, PM, the major component of AirP, exists in three categories: PM_10_ (coarse, <10µm), PM_2.5_ (fine, <2.5µm), and PM_0.1_ (ultrafine, <1µm). PM_2.5_ and PM_0.1_ exhibit heightened cytotoxicity due to their deeper penetrance into the lungs and nasal passageways^64^. DEP particles are predominantly ultrafine (PM_0.1,_-PM_2.5_), while WTC is largely coarse (PM_10_)^51,65^. Despite these differences, both DEP and WTC contained abundant transition metals, including Fe, Cu, Zn, Mn, and post-transition metal Pb (**S.Fig. 1C**), that are capable of catalyzing lipid peroxidation and implicated in amyloidogenic processing. Thus, while DEP and WTC dust differ fundamentally in their ionic and carbonaceous profiles, their shared metal enrichment provides the most plausible mechanistic link between these distinct exposures and AD pathology. In contrast to DEP and WTC, WS contained at least 1,000-fold lower metal content (**S.Fig 1D**). Here, we compared the DEP and WTC exposures to acute WS exposure, which was performed separately and at a higher concentration, exclusively in female mice. Due to this asymmetry, WS results are presented in the supplemental figures.

### DEP and WTC elicit convergent transcriptomic responses despite chemical heterogeneity

Acute exposure to DEP or WTC induced robust transcriptional responses in the cortex, a region moderately afflicted by AD neuropathology. DEP exposure altered over 4,000 genes, while WTC exposure altered approximately 1,800 genes, with only ∼200 genes differing between the two exposures, suggesting high overlap in their transcriptomic response (**Fig. 1A-C**). GSEA revealed shared upregulation of oxidative stress, IFNα, and DNA repair pathways, alongside downregulation of protein secretion and UV response pathways compared to filtered air (**Fig. 1D,E**).

**Figure 1:**
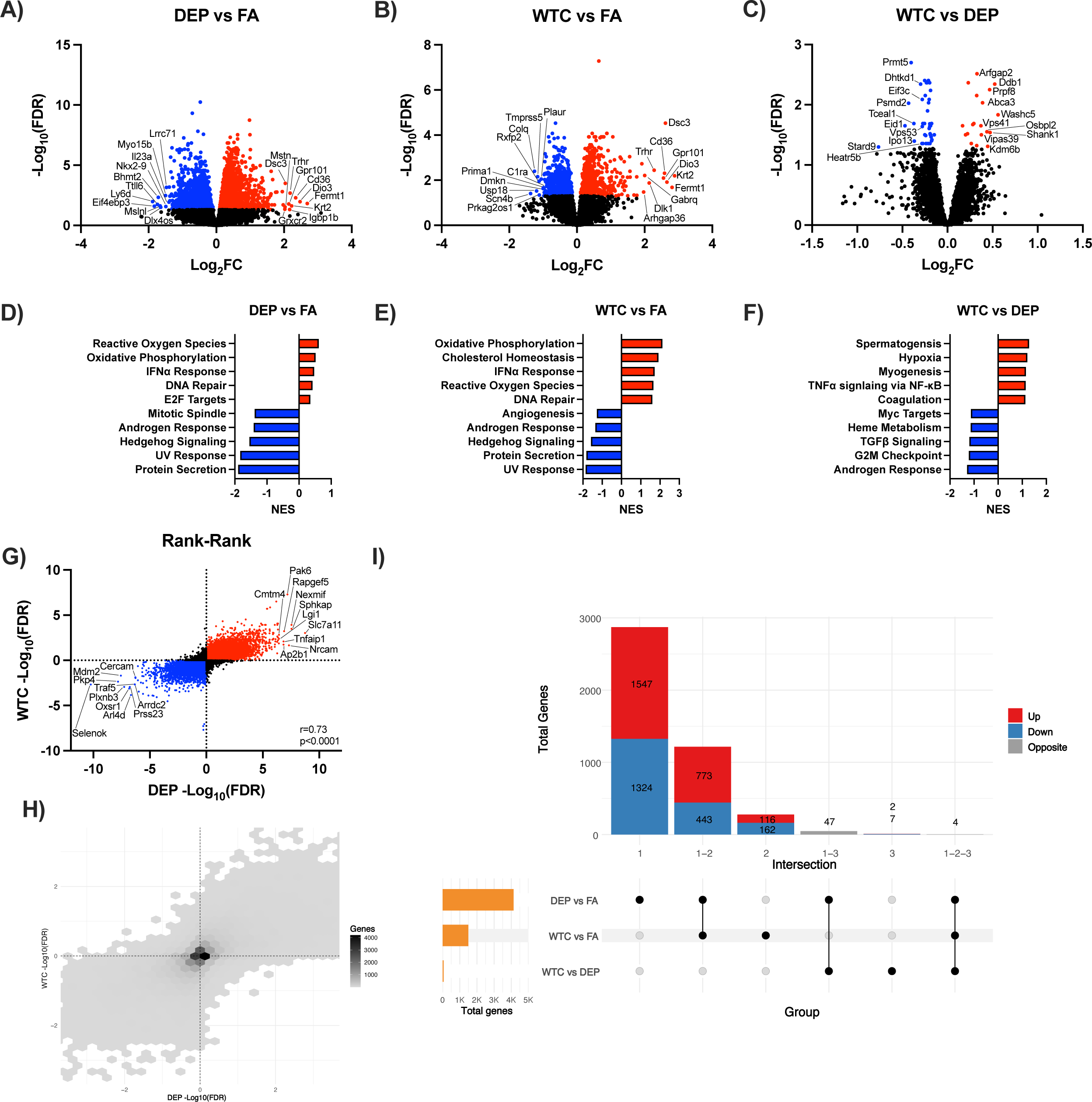
Acute 5-hour exposure to DEP or WTC results in highly conserved transcriptomic responses. Volcano plots of genes for **A)** DEP, **B)** WTC, from FA, or **C)** between WTC and DEP. GSEA pathway enrichment for **D)** DEP, **E)** WTC, from FA or **F)** between WTC and DEP. **G)** Rank-rank plot comparing all genes for DEP and WTC, and **H)** a rank-rank hex plot for gene density. **I)** Upset plot showing the number of DEGs exclusive or shared between the exposures in the same direction of change.

DEP exposure preferentially enhanced NF-κB signaling, whereas WTC exposure was associated with increased heme metabolism, suggestive of cerebral microhemorrhages (**Fig. 1F**). Importantly, Ingenuity Pathway Analysis predicted activation of amyloid precursor protein (APP) and microtubule associated protein tau (MAPT) regulatory networks following both exposures, directly linking these convergent transcriptional responses to AD-relevant pathways.

Rank-rank correlation analysis demonstrated a strong concordance between DEP- and WTC-induced gene expression changes (r=0.73; **Fig. 1G**), indicating that despite stark differences in chemical composition and particle size, both exposures elicit common gene signatures. Hexbin visualization further confirmed extensive overlap among all genes across both exposures (**Fig. 1H**). Consistent with these findings, upset plot analysis demonstrated that approximately 1,200 genes were regulated in the same direction by both exposures (intersection 1–2), whereas only 47 genes showed discordant regulation (intersection 1–3; **Fig. 1I**). DEG analysis was split by sex in **Supplemental Figure 2**. Collectively, these findings demonstrate that chemically distinct, but metal-rich AirP sources converge on a shared cortical stress transcriptome, characterized by oxidative stress, interferon signaling, and altered protein trafficking.

### WS exposure elicits a distinct transcriptomic response that does not overlap with DEP or WTC

In contrast to DEP and WTC, WS exposure did not share overlapping top differentially expressed genes (DEGs) with either metal-rich exposure (**S.Fig. 3A**). Although WS modestly engaged IFNα signaling, it uniquely enriched IFNγ signaling, fatty acid metabolism, and unfolded protein response pathways (**S.Fig. 3B**). Rank-rank correlation and hexbin analyses revealed no meaningful concordance between WS and either DEP or WTC (**S.Fig. 3C-F**). Upset plots of the three exposures showed few DEGs with high discordance (Intersection 1-3, **S.Fig. 3G**). These data indicate that the convergent transcriptomic response observed following DEP and WTC exposure is not a generic consequence of particulate exposure, but rather reflects a shared biological response associated with metal-rich AirP.

### DEP and WTC exposures induce ferroptotic priming by increased lipid peroxidation and blunted antioxidant defense

Given the shared oxidative stress signature induced by DEP and WTC, we examined whether these transcriptomic changes translated into ferroptosis-relevant biochemical alterations, particularly in the antioxidant systems responsible for detoxifying lipid peroxides and maintaining glutathione (GSH) homeostasis (**Fig. 2A**). Transcriptomic analyses revealed coordinated dysregulation of ferroptosis-associated genes, including reduced expression of glutathione peroxidase 4 (GPx4) and glutathione synthetase (GSS), alongside increased expression of pro-ferroptotic mediators such as PTGS2 and ACSL4 (**Fig. 2B**). In parallel, the iron-responsive surface receptor CD44 and the cystine importer SLC7A11, critical for cystine uptake and GSH synthesis, were upregulated, consistent with a compensatory response to oxidative stress and increased iron presence (**Fig. 2B**). Consistent with these changes, lipid peroxidation increased following both exposures, with 4-hydroxynonenal (HNE) levels rising by at least 30% following DEP and WTC exposure (**Fig. 2C**). In contrast, protein nitration product, nitrotyrosine (NT) protein levels, and HNE assessed by immunofluorescence (IF) remained unchanged at this acute time point (**S.Fig. 4A,B**), indicating early lipid-selective oxidative injury.

**Figure 2:**
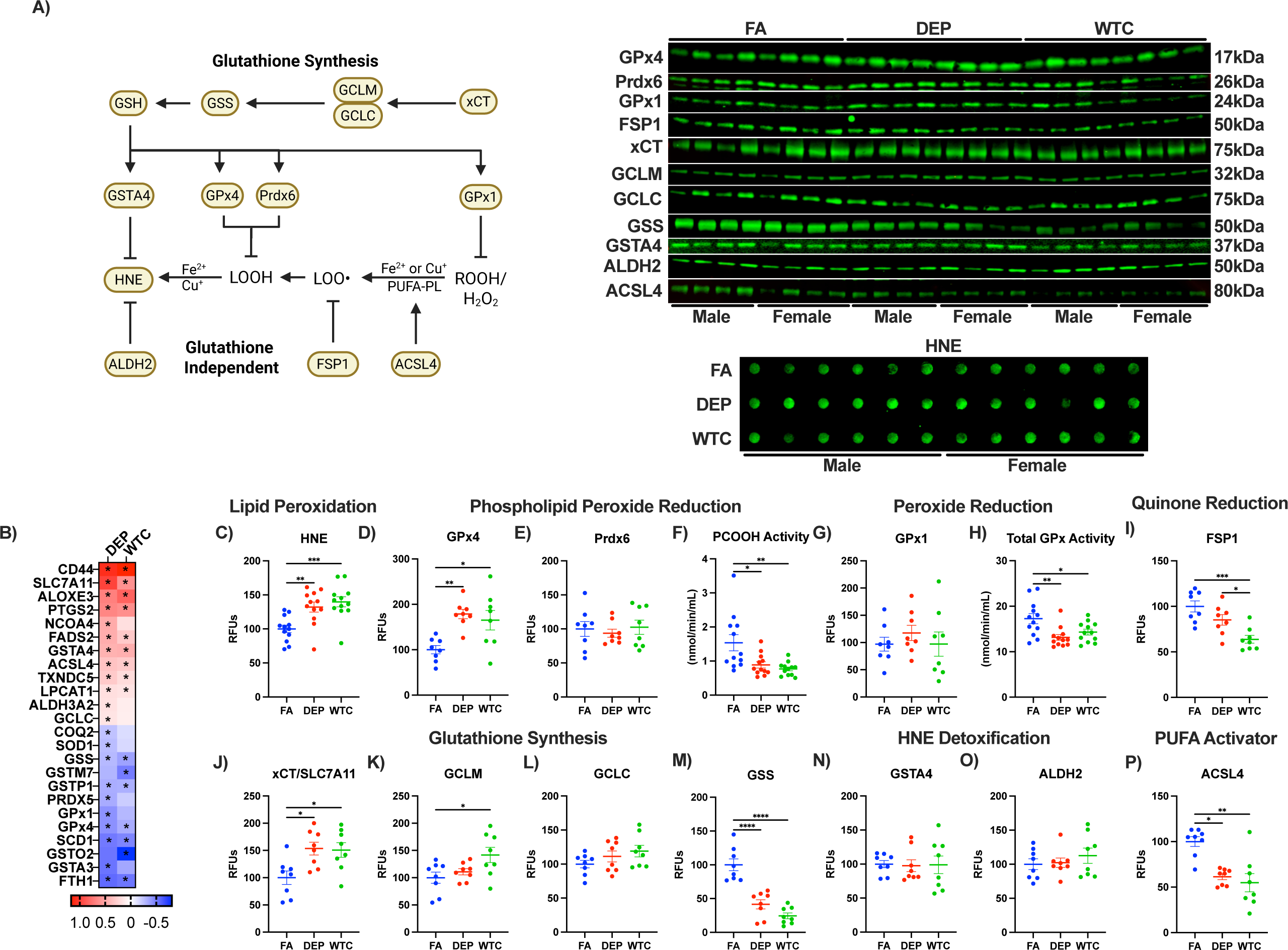
Acute air pollution exposure sensitizes the brain to ferroptotic changes. **A)** Schematic representation of glutathione-dependent and glutathione-independent antioxidant pathways regulating lipid peroxide detoxification. **B)** DEGs related to ferroptosis in the frontal cortex exposed to DEP and WTC for 5-hours. Cortical levels of **C)** HNE, **D)** GPx4, **E)** Prdx6, **F)** PCOOH activity, **G)** GPx1, **H)** total GPx activity, **I)** FSP1, **J)** SLC7A11, **K)** GCLM, **L)** GCLC, **M)** GSS, **N)** GSTA4, **O)** ALDH2, and **P)** ACSL4 measured in whole cell lysates by Western blot, dot blot, or enzymatic assay. Statistical analysis was performed using ANCOVA adjusted for sex with Bonferroni’s post hoc test. *p<0.05, **p<0.01, ***p<0.001, ****p<0.0001.

We found GPx4 protein abundance to increase with both DEP and WTC (**Fig. 2D**), while Prdx6 and Gpx1 protein levels remained unchanged (**Fig. 2E, G**). Functional assays revealed a ∼50% reduction in phospholipid hydroperoxide detoxification capacity following both DEP and WTC exposure (**Fig. 2F**), accompanied by reduced total GPx activity (**Fig. 2H**), indicating a selective deficit in lipid peroxide detoxification rather than global peroxide handling. To further evaluate ferroptosis-relevant antioxidant systems, we examined ferroptosis suppressor protein 1 (FSP1), which limits lipid peroxidation through quinone reduction. FSP1 protein levels were reduced by 35% with DEP relative to FA and by 25% relative to DEP exposure following WTC exposure (**Fig. 2I**). These deficits occurred alongside impaired glutathione synthesis, as GSS protein levels declined by >60% despite compensatory increases in cystine transport and upstream GSH pathway components (**Fig. 2J–M**).

Proteins involved in the detoxification of lipid-derived aldehydes, including glutathione S-transferase A4 (GSTA4) and aldehyde dehydrogenase 2 (ALDH2), were unchanged following DEP and WTC exposure (**Fig. 2N,O**). Notably, ACSL4 protein levels declined by at least 40% following both exposures (**Fig. 2P**), suggesting disruption of phospholipid remodeling pathways that influence ferroptotic susceptibility. Collectively, these findings indicate that acute DEP and WTC exposure induces a state of ferroptotic priming characterized by lipid peroxide accumulation, impaired detoxification capacity, and compromised glutathione synthesis, rather than overt ferroptotic cell death.

### WS exposure does not induce ferroptotic priming despite altered peroxide-handling capacity

In contrast to DEP and WTC, WS exposure did not increase levels of HNE or NT (**S.Fig. 4C,D**), indicating an absence of overt lipid or protein oxidative damage at this time point. GPx4, Prdx6, and GPx1 protein levels were unchanged with WS exposure (**S.Fig. 4E,F,H**). Although phospholipid peroxide detoxification activity was reduced by 80% following WS exposure (**S.Fig. 4G**), total GPx activity increased by 30% (**S.Fig. 4I**), suggesting compensatory engagement of non-lipid peroxide detoxification pathways. WS exposure did not alter most ferroptosis-related proteins aside from a reduction in ALDH2 (**S.Fig. 4J-P**). These findings indicate that WS exposure lacks the coordinated ferroptotic signature observed following metal-rich AirP.

### Copper handling pathways are differentially regulated by DEP, WTC, and WS exposures

Given emerging links between copper dysregulation, oxidative stress, and neurodegeneration, we examined copper handling pathways following acute AirP exposure. Copper uptake is mediated by the transporter CTR1 and buffered by metallothioneins, while cytosolic chaperones such as copper chaperone for superoxide dismutase (CCS) and ATOX1 deliver copper to antioxidant and mitochondrial targets, including cytochrome c oxidase via COX17 and COA6 (**S.Fig. 4Q**). At the transcriptional level, multiple copper-handling genes were downregulated following DEP exposure and showed similar directional trends following WTC exposure, whereas WS exposure produced opposing expression patterns (**S.Fig. 4R**). These findings suggest selective remodeling of copper buffering and trafficking pathways by metal-rich AirP.

At the protein level, CCS abundance was reduced by at least 35% following WTC exposure relative to both filtered air and DEP (**S.Fig. 4S**). In contrast, SOD1 protein abundance and total superoxide dismutase activity were unchanged following DEP or WTC exposure (**S.Fig. 4T,U**), indicating preserved basal copper-dependent antioxidant capacity. WS exposure did not alter SOD1 protein levels but reduced total SOD activity by approximately 50% (**S.Fig. 4V,W),** consistent with functional impairment independent of SOD1 abundance. Together, these data indicate that acute metal-rich AirP exposure perturbs copper trafficking and buffering without engaging overt cuproptosis or SOD1-dependent oxidative collapse, whereas WS disrupts copper-dependent antioxidant activity through a distinct mechanism.

### DNA damage but not transcript errors accompany oxidative stress induced by DEP and WTC

To determine whether oxidative stress induced by metal-rich AirP extended beyond lipid damage, we examined markers of nucleic acid integrity and transcript surveillance. Transcriptomic analysis revealed coordinated regulation of genes involved in DNA damage signaling and repair following DEP exposure, with similar but weaker trends observed following WTC exposure (**S.Fig. 5A**). These included regulators of DNA damage sensing (ATM, CHEK1, GADD45A), base excision repair (APEX1, NEIL1, POLB), and nucleotide excision repair (XPA, ERCC6L2), consistent with activation of oxidative DNA damage response pathways.

Consistent with these transcriptional changes, IF revealed increased cortical levels of 8-hydroxy-2′-deoxyguanosine (8-OHdG) following DEP exposure, whereas WTC exposure did not significantly alter 8-OHdG at this acute time point (**S.Fig. 5B**). These findings indicate that DEP induces greater nucleic acid oxidation than WTC, paralleling the more pronounced lipid peroxidation and antioxidant impairment observed following DEP exposure.

To assess whether oxidative stress extended to transcript integrity, we performed a pilot Circle-sequencing (CircSeq) analysis to detect transcript errors. In this limited dataset (male mice; n=1 per group), no consistent or exposure-specific increase in transcript errors was detected across transcripts generated by RNA polymerase I, II, III, or mitochondrial RNA polymerase (**S.Fig. 5C-F**). Given the absence of a clear signal and the exploratory nature of this analysis, further CircSeq experiments were not pursued.

Collectively, these findings demonstrate that acute DEP exposure induces DNA oxidation and engages DNA repair pathways without causing notable transcriptional mutagenesis, consistent with secondary nucleic acid stress arising downstream of oxidative injury rather than global transcriptional infidelity.

### Acute AirP exposure alters iron handling and heme metabolism without increasing total brain iron

To determine whether the ferroptosis-relevant oxidative imbalance observed following DEP and WTC exposure was associated with changes in brain iron availability or handling, we examined proteins involved in iron transport, storage, export, and heme metabolism (**Fig. 3A**). Iron primarily enters the cell bound to transferrin (TF) via the transferrin receptor (TfR) as Fe^3+^ or through divalent metal transporter 1 (DMT1) as Fe^2+^. Intracellular iron is stored within ferritin complexes composed of ferritin heavy (FTH1) and light (FTL) chains, while iron export occurs exclusively through ferroportin (FPN) or through NCOA4-mediated ferritinophagy.

**Figure 3:**
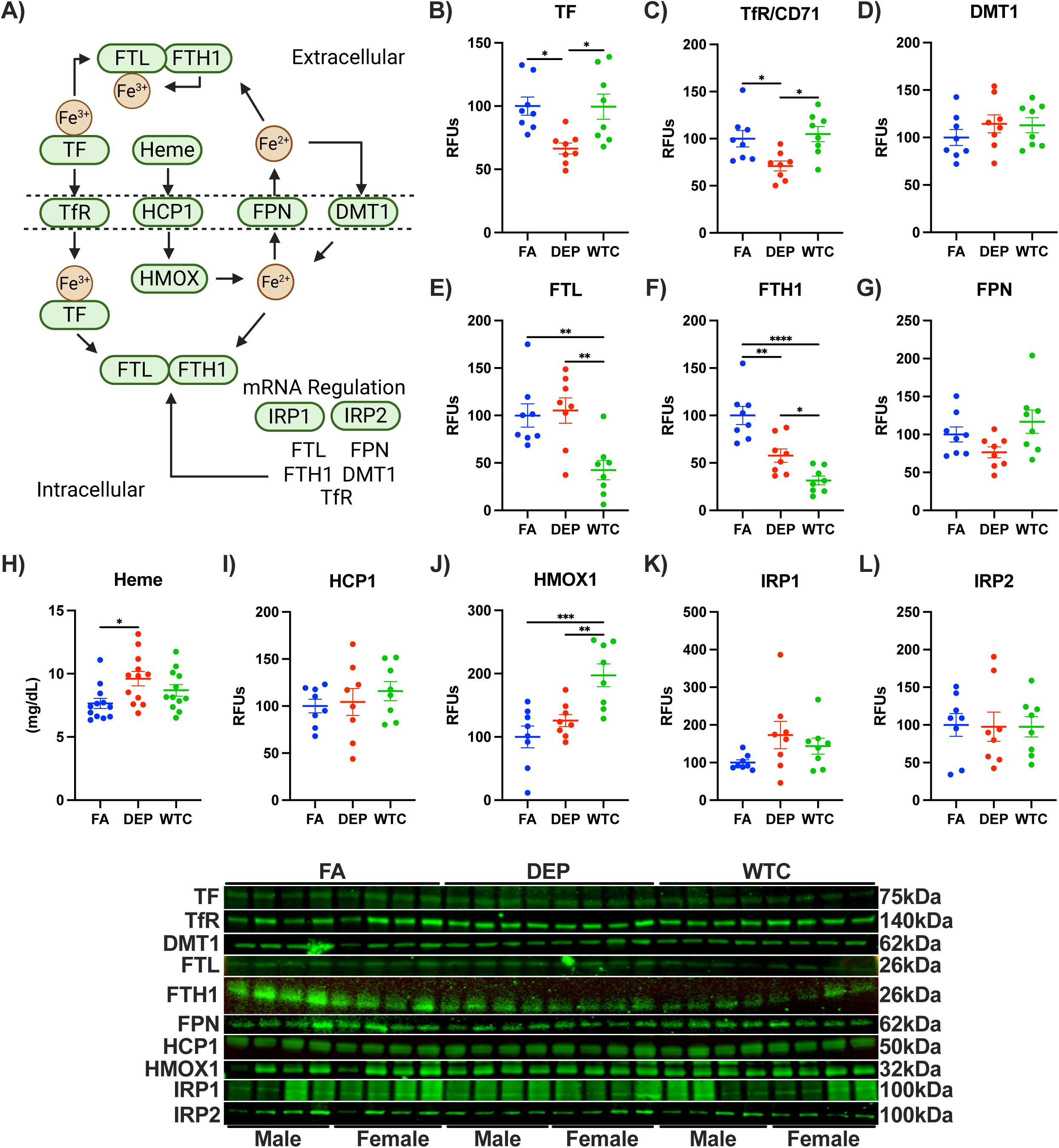
Iron signaling is downregulated in response to acute air pollution exposure. **A)** Schematic representation of iron signaling pathways. Western blots or assay for **B)** TF, **C)** TfR, **D)** DMT1, **E)** FTL, **F)** FTH1, **G)** tissue heme, **H)** HCP1, **I)** HMOX1, **J)** FPN, **K)** IRP1, and **L)** IRP2. Statistical analysis was performed using ANCOVA adjusted for sex with Bonferroni’s post hoc test. *p<0.05, **p<0.01, ***p<0.001, ****p<0.0001.

Despite evidence of ferroptotic priming, MRI-based measurements revealed no changes in total brain iron content following DEP or WTC exposure (**S.Fig. 6A,B**), indicating that acute exposure does not produce bulk iron accumulation. Instead, both exposures altered iron handling pathways. DEP reduced TF and TfR abundance, whereas WTC primarily impaired ferritin-based iron storage, with pronounced reductions in FTH1 and FTL (**Fig. 3B-F**). DMT1 and FPN1 protein abundance were unchanged following either DEP or WTC exposure (**Fig. 3D,G**), suggesting selective suppression of transferrin-dependent iron uptake rather than generalized iron import or export. Overall, these changes indicate impaired ferritin-based iron sequestration, particularly following WTC exposure.

Consistent with altered iron storage, tissue heme levels increased by 25% following DEP exposure and trended upward with WTC exposure compared to FA (**Fig. 3H**). Despite increased heme content, expression of the heme importer HCP1 was unchanged (**Fig. 3I**). In contrast, heme oxygenase 1 (HMOX1), which catalyzes heme degradation and liberates Fe^2+^, increased twofold following WTC exposure (**Fig. 3J**), suggesting enhanced heme turnover and potential microvascular stress. In contrast, canonical iron-sensing pathways remained intact (**Fig. 3K,L**). These data indicate that acute metal-rich AirP redistributes iron toward bioactive pools without increasing bulk iron accumulation.

### WS exposure does not induce comparable alterations in iron handling

Consistent with its low metal content, WS exposure did not induce iron redistribution or heme metabolism changes observed with metal-rich AirP. Instead, modest alterations in iron regulatory proteins were observed without changes in iron storage or export (**S.Fig. 6C-M).** Collectively, these findings indicate that WS exposure does not induce the iron redistribution and heme-associated responses observed with the metal-rich DEP and WTC.

### Acute DEP and WTC exposure enhances amyloidogenic processing and blunts lipid raft antioxidant defenses

We previously identified LRs as hotspots for lipid peroxidation^26,27^, which may enhance amyloid processing^35,66^ (**Fig. 4A**). Given the sensitivity of lipid rafts (LRs) to lipid peroxidation and their central role in APP processing, we examined amyloid-related pathways. DEP exposure broadly upregulated transcripts involved in APP processing (ADAM10, BACE1, PSEN1, APH1B), endosomal trafficking (VPS35, VPS26B, SNX27, SORT1), and tau kinase signaling (CDK5R1, GSK3β, MARK1, TTBK2), whereas WTC produced a more restricted transcriptional response. Notably, ABCA7 was the only transcript significantly downregulated by both exposures (**Fig. 4B**). These findings highlight impaired lipid and amyloid handling as a shared vulnerability, while uniquely engaging DEP in broader amyloidogenic and tau-related genes.

**Figure 4:**
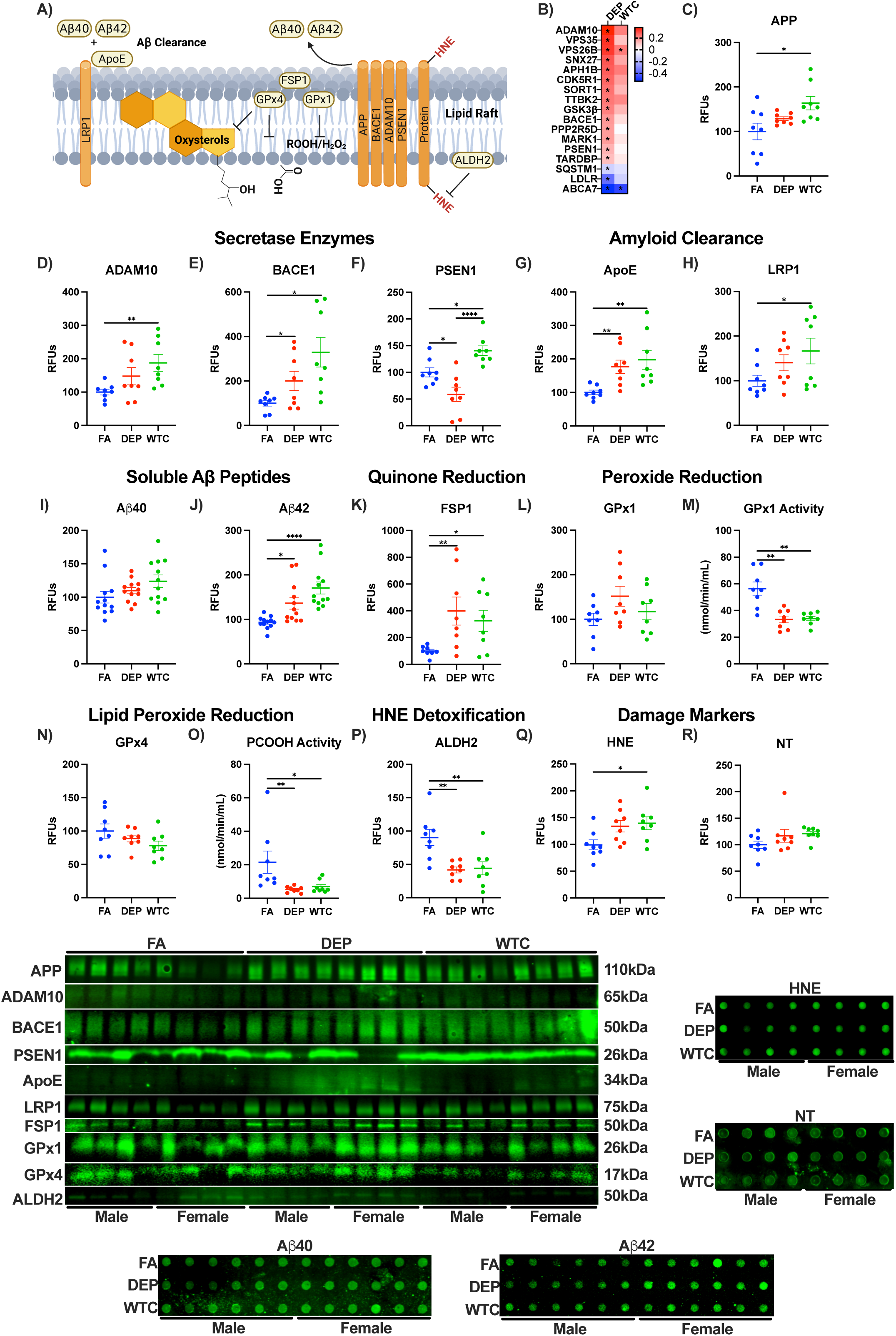
Amyloidogenic processing and damage to the lipid raft increase with acute air pollution exposure. **A)** Schematic representation integrating lipid raft antioxidant defense, damage, and APP processing. **B)** heatmap of DEGs related to AD neuropathology. Enzymatic activities, proteins by Western blot or damage markers by dotblot in lipid rafts or soluble Aβ peptides by dotblot: **C)** APP, **D)** ADAM10, **E)** BACE1, **F)** PSEN1, **G)** ApoE, **H)** LRP1, **I)** Aβ40, **J)** Aβ42, **K)** FSP1, **L)** GPx1, **M)** GPx1 activity, **N)** GPx4, **O)** PCOOH activity, **P)** ALDH2, **Q)** HNE, and **R)** NT. Statistical analysis was performed using ANCOVA adjusted for sex with Bonferroni’s post hoc test. *p<0.05, **p<0.01, ***p<0.001.

Consistent with these transcriptional changes, protein-level analysis demonstrated exposure-specific alterations in APP processing components. LR APP protein levels increased by 60% following WTC exposure but were unchanged following DEP exposure (**Fig. 4C**). The α-secretase ADAM10 increased two-fold following WTC exposure and trended upward with DEP (**Fig. 4D**). In contrast, the β-secretase BACE1 increased two-fold with DEP exposure and three-fold with WTC exposure (**Fig. 4E**). The catalytic γ-secretase subunit presenilin-1 (PSEN1) decreased by 40% following DEP exposure but increased by 40% following WTC exposure (**Fig. 4F**), indicating divergent regulation of secretase components, but converged on increased amyloidogenic output.

Apolipoprotein E (ApoE), which binds amyloid peptides and facilitates their clearance, increased twofold following both DEP and WTC exposure (**Fig. 4G**). LR protein of the ApoE receptor, LRP1, increased by 65% following WTC but was unchanged following DEP (**Fig. 4H**). Measurement of soluble amyloid peptides in whole cell lysates revealed trends of increase in Aβ40 following both exposures, whereas the more aggregation-prone Aβ42 increased by 35% following DEP exposure and by 70% following WTC exposure (**Fig. 4I,J**), indicating a shift in amyloidogenic production.

To determine whether altered amyloid processing occurred in parallel with compromised LR antioxidant defenses, we examined LR-specific protective mechanisms and damage. FSP1, increased fourfold following DEP and threefold following WTC exposures (**Fig. 4K**), consistent with a compensatory response to oxidative stress. Within lipid rafts, GPx1 protein abundance was unchanged, but GPx1 enzymatic activity declined by 40% following both DEP and WTC exposures (**Fig. 4L,M**). GPx4 protein trended downward in LRs following both exposures, while phospholipid hydroperoxide (PCOOH) reduction capacity decreased by at least 70% (**Fig. 4N,O**), indicating impairment of lipid peroxide detoxification within the LR. In parallel, LR ALDH2 levels decreased by at least 55% following both DEP and WTC exposure, while LR HNE, but not LR NT, increased by 40% following WTC exposure and trended upward with DEP (**Fig. 4P-R**). Collectively, these findings demonstrate that acute exposure to metal-rich AirP enhances amyloidogenic processing and promotes accumulation of the aggregate-prone Aβ42 while simultaneously impairing lipid raft antioxidant defenses, recapitulating biochemical features observed in postmortem Alzheimer’s disease brain tissue^26,27,54^.

### WS exposure alters amyloid peptide profiles without inducing the amyloidogenic signature observed with DEP or WTC

To determine whether WS exposure influenced amyloid pathways, we examined APP processing and amyloid peptide production in whole cell lysates. In contrast to DEP and WTC, WS exposure reduced APP protein abundance by 40% (**S.Fig. 7A**). WS did not alter ADAM10 or BACE1 levels, but reduced PSEN1 levels by 50% (**S.Fig. 7B-D**). Despite these changes, WS exposure increased soluble Aβ40 by 85%, without altering Aβ42 levels (**S.Fig. 7E,F**), indicating a shift toward production of the less aggregation-prone amyloid species. ApoE levels increased by 40%, while LRP1 abundance was unchanged (**S.Fig. 7G,H**). These data demonstrate that WS exposure alters amyloid peptide balance without engaging the amyloidogenic machinery or lipid raft dysfunction observed following metal-rich AirP, further reinforcing mechanistic divergence between WS and DEP/WTC.

### Acute AirP exposure engages xenobiotic, redox, and other stress-adaptive transcriptional responses

To define regulatory programs engaged by acute air pollution exposure, we examined transcription factors associated with xenobiotic metabolism, oxidative stress, and inflammatory signaling (**Fig. 5A**). Transcriptomic analysis revealed increased expression of MEF2C, a regulator of neuronal development and synaptic function, following both DEP and WTC exposure, alongside DEP-specific induction of the aryl hydrocarbon receptor (AhR) and its neuronal binding partner ARNT2 (**Fig. 5B**), which mediate transcriptional responses to polycyclic aromatic hydrocarbons (PAHs) and related xenobiotics. In parallel, transcripts encoding the AhR repressor (AhRR), ARNT1, and ATF4 were reduced, consistent with engagement of xenobiotic-responsive transcriptional circuits (**Fig. 5B**).

**Figure 5:**
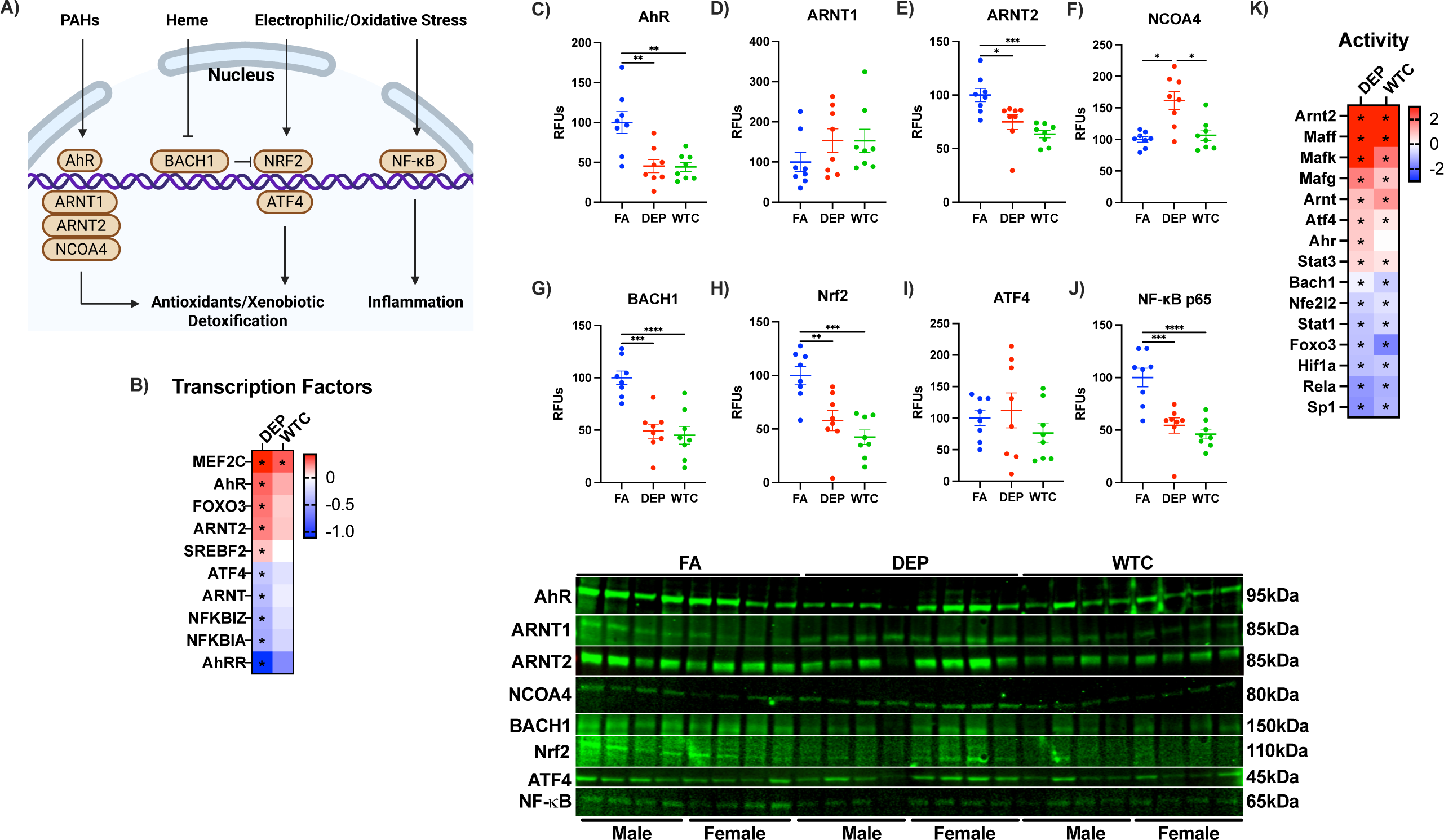
Transcription factors involved in response to air pollution, oxidative stress, and inflammation. **A)** Schematic representation of nuclear stress-sensing pathways linking xenobiotic exposure, oxidative stress, and inflammatory signaling. **B)** DEGs for transcription factors found in RNA-seq. Western blots of nuclear lysates for **C)** AhR, **D)** ARNT1, **E)** ARNT2, **F)** NF-κB, **G)** Nrf2, **H)** ATF4, and **I)** NCOA4. **J)** DoRothEA derived transcription factor activity from RNA-seq. Statistical analysis was performed using ANCOVA adjusted for sex with Bonferroni’s post hoc test. *p<0.05, **p<0.01, ***p<0.001, ****p<0.0001.

Assessment of nuclear protein abundance revealed partial divergence from transcript-level changes, consistent with the temporal dynamics of transcription factor activation. Nuclear AhR protein levels were reduced following both DEP and WTC exposure, while ARNT1 abundance was unchanged (**Fig. 5C,D**). Neuronal ARNT2 levels declined following both exposures, whereas nuclear NCOA4 increased selectively following DEP exposure (**Fig. 5E,F**), suggesting exposure-specific modulation of xenobiotic and iron transcriptional responses.

To evaluate redox-associated transcriptional control, we assessed BACH1 and Nrf2, which competitively regulate antioxidant and iron metabolism genes. BACH1 protein levels declined by at least 50% following both DEP and WTC exposure (**Fig. 5G**), consistent with elevated heme levels and relief of BACH1-mediated repression. In contrast, nuclear Nrf2 levels declined by 45% following DEP exposure and 60% following WTC exposure, while ATF4 protein abundance remained unchanged (**Fig. 5H,I**), suggesting redistribution rather than amplification of canonical antioxidant signaling. Nuclear NF-κB p65 levels were also significantly reduced following DEP and WTC exposure (**Fig. 5J**), indicating selective remodeling rather than global inflammatory activation.

Because transcription factor activity is not fully captured by steady-state nuclear abundance, we inferred regulatory activity using DoRothEA based on validated target gene expression (**Fig. 5K**). This analysis revealed increased activity of AhR pathway components, including AhR itself, following DEP exposure, and its obligate partners ARNT and ARNT2 following both DEP and WTC exposure. In parallel, activity of small Maf proteins (MAFF, MAFK, MAFG), which heterodimerize with Nrf2 to regulate antioxidant and detoxification genes, and ATF4 were increased, despite reduced Nrf2 (NFE2L2) activity. AirP exposure also selectively remodeled inflammatory and stress signaling pathways. STAT3 activity increased, whereas RELA (NF-κB p65) and STAT1 activity declined, indicating noncanonical inflammatory regulation. Transcription factors associated with cellular resilience and metabolic adaptation, including FOXO3, HIF1A, and SP1, were reduced following DEP and WTC exposure. Together, these data indicate that acute metal-rich AirP exposure engages coordinated xenobiotic metabolism, redox modulation, and stress-adaptive transcriptional programs without global inflammatory activation, consistent with an early adaptive response to oxidative challenge.

### WS exposure induces limited and divergent transcription factor responses

In contrast to DEP and WTC, WS exposure elicited a more restricted and divergent transcription factor response. Nuclear AhR levels were unchanged, while ARNT1 and ARNT2 abundance declined (**S.Fig. 8A-C**). NCOA4 increased 2.5-fold following WS exposure (**S.Fig. 8D**), suggesting limited engagement of xenobiotic response signaling. Unlike metal-rich AirP, WS increased nuclear BACH1 levels while reducing Nrf2 (**S.Fig. 8E,F**), indicating a shift toward transcriptional repression of antioxidant and iron-handling genes. ATF4 protein abundance increased, while NF-κB p65 levels were unchanged (**S.Fig. 8G,H**). Consistent with these findings, WS exposure produced minimal transcription factor activity changes by DoRothEA analysis, further supporting the absence of coordinated xenobiotic or redox regulatory engagement.

### DEP and WTC exposures converge on shared neuronal and oligodendrocyte transcriptional responses

To determine whether bulk transcriptomic changes reflected intrinsic transcriptional regulation or shifts in cellular composition, cell-type deconvolution was performed on our RNA-seq dataset. Both DEP and WTC exposures were associated with a reduction in oligodendrocyte proportions and a reciprocal increase in neuronal proportions relative to FA (**Fig. 6A,B**). Because changes in cellular composition can mask or distort DEGs, we next adjusted the bulk RNA-seq analysis for oligodendrocyte and neuronal proportions. After adjusting for cellular composition, additional ferroptosis-, lipid remodeling-, and vascular-associated genes emerged (**Fig. 6C-K**) in DEP-exposed brains. In contrast, WTC exposure revealed population-adjusted changes associated with heme metabolism and synaptic modulation (**Fig. 6L,M**).

**Figure 6:**
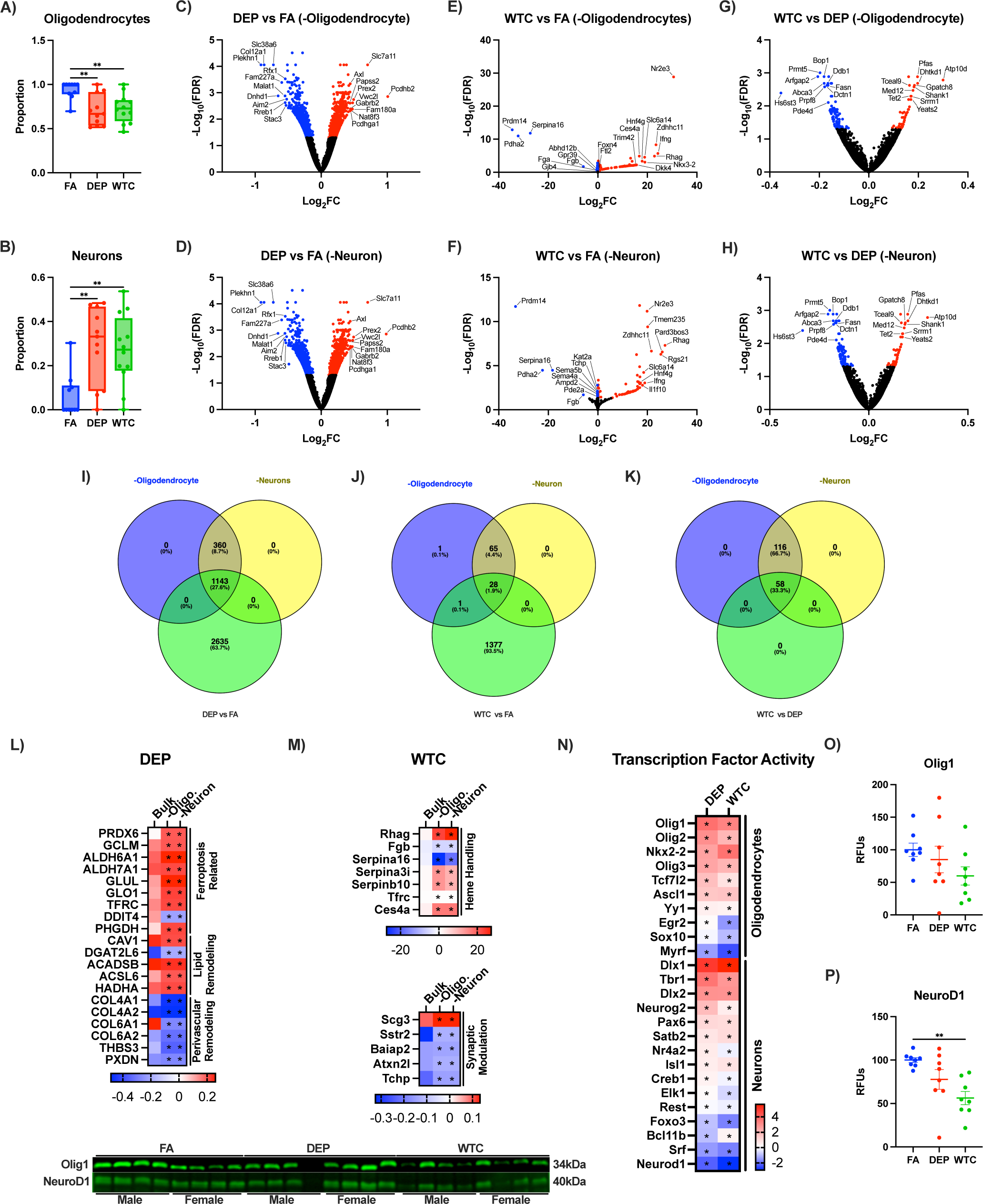
Oligodendrocyte and neuron populations are altered with acute air pollution exposure. **A)** Deconvolve analysis of oligodendrocytes and oligodendrocyte adjusted volcano plots for **B)** DEP vs FA, **C)** WTC vs FA, and **D)** WTC vs DEP. **E)** Deconvolve analysis of neurons and neuron-adjusted volcano plots for **F)** DEP vs FA, **G)** WTC vs FA, and **H)** WTC vs DEP. Venn diagrams highlighting DEGs exclusive to cell type adjusted analysis for **I)** DEP vs FA, **J)** WTC vs FA, and **K)** WTC vs DEP. Heatmaps of genes related to ferroptosis for **L)** DEP, **M)** WTC, and **N)** transcription factor activity by DoRothEA. Western blots of nuclear lysate for **O)** Olig1 and **P)** NeuroD1. Statistical analysis was performed using ANOVA with Kruskal-Wallis for A,E, and ANCOVA adjusted for sex with Bonferroni’s post hoc test for O,P. **p<0.01

To further assess whether these changes reflected engagement of lineage-associated transcriptional responses rather than cell loss or gain alone, transcription factor activity was again inferred using DoRothEA. Transcription factor inference revealed increased activity of oligodendrocyte lineage regulators and reduced neuronal transcriptional programs, without evidence of overt cell loss (**Fig. 6N**). At the protein level, nuclear Olig1 abundance was unchanged following DEP or WTC exposure, whereas NeuroD1 protein levels were reduced by 45% following WTC exposure, mirroring the DoRothEA activity (**Fig. 6O,P**). Consistent with the absence of overt cell loss, levels of myelin basic protein (MBP) and the neuronal marker NeuN were unchanged following DEP and WTC exposure (**S.Fig. 9A,B**).

### WS exposure shifts cell-state responses in the opposite direction of metal-rich AirP

Deconvolution analysis of WS exposure revealed a contrasting pattern, characterized by increased oligodendrocyte proportions and reduced neuronal proportions (**S.Fig. 9C,D**). MBP protein levels were unchanged, while NeuN levels decreased by 40% following WS exposure (**S.Fig. 9E,F**). In contrast to DEP and WTC, WS exposure produced distinct transcription factor responses, with increased activity of oligodendrocyte-associated transcription factors including TCF12, PRDM14, SOX11, KLF9, MEIS1, and EBF1, accompanied by reduced activity of the neuronal transcription factor FOXP2 (**S.Fig. 9G**). NeuroD1 was not identified in DoRothEA analysis and was unchanged at the protein level following WS exposure (**S.Fig. 9H**). Collectively, these findings demonstrate that DEP and WTC exposure converge on shared neuronal and oligodendrocyte transcriptional responses that persist after accounting for cellular composition, whereas WS exposure induces a divergent cell-state response characterized by oligodendrocyte enrichment and reduced neuronal representation.

### DEP and WTC exposure alters white-matter microstructure and induces lipid peroxidation in the corpus callosum

To determine whether acute AirP exposure alters brain microstructure, *ex vivo* diffusion tensor imaging (DTI) was performed across multiple brain regions. DTI revealed increased fractional anisotropy (**Fig. 7A**) and reduced diffusivity (**Fig. 7B**) in the corpus callosum following DEP and WTC exposure, and these findings were confined to white matter. Other brain regions exhibited minimal or no changes in these measurements. These observations are consistent with acute alterations in white-matter microstructure, suggestive of increased directional restriction of water diffusion, potentially reflecting cellular or myelin-associated swelling rather than tissue loss.

**Figure 7:**
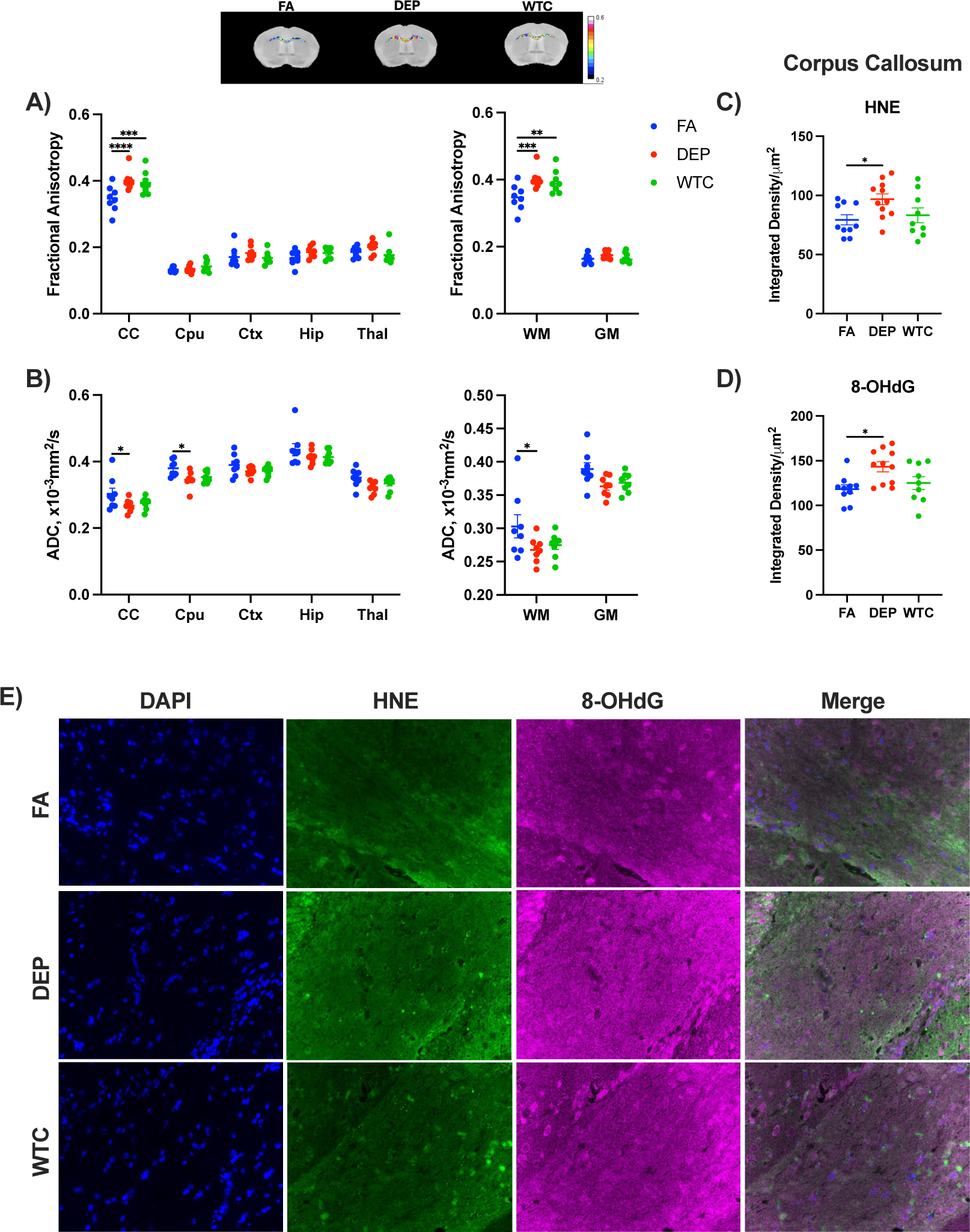
Structural changes in the Corpus Callosum consistent with white matter damage. **A)** Fractional Anisotropy and **B)** Apparent diffusion coefficient (ADC) measured by brain region and white and grey matter by *ex vivo* MRI. Corpus callosum for **C)** HNE, **D)** 8-OHdG, and **E)** representative images for immunofluorescent microscopy at 40x. Statistical analysis was performed using two-way ANOVA for A,B, or ANCOVA adjusted for sex with Bonferroni’s post hoc test for C,D. *p<0.05, **p<0.01, ***p<0.001, ****p<0.0001.

DEP exposure uniquely increased lipid peroxidation and DNA damage within the corpus callosum (**Fig. 7C,D**), whereas WTC altered diffusion properties without overt oxidative injury. These findings demonstrate that acute metal-rich AirP rapidly perturbs white-matter integrity, with DEP producing the strongest oxidative injury. Collectively, these data demonstrate that acute AirP exposure rapidly perturbs white-matter microstructure, with the corpus callosum exhibiting the greatest sensitivity. The dissociation between diffusion abnormalities and oxidative damage across exposures suggests that microstructural alterations can arise in the absence of overt lipid or DNA oxidation.

### WS exposure does not induce oxidative damage or inflammation in the white matter or hippocampus

To determine whether acute WS exposure elicited oxidative or inflammatory injury within white matter or hippocampal regions, IF was performed in the corpus callosum and across multiple hippocampal subregions. In the corpus callosum, WS exposure did not alter levels of HNE, 8-OHdG, degraded myelin basic protein (dMBP), the complement component C5 or its activation fragment C5a, or the microglial marker Iba1 (**S.Fig. 10A-F**). Similarly, WS exposure did not alter HNE, 8-OHdG, dMBP, C5, C5a, or Iba1 immunoreactivity within hippocampal subregions, aside from increased dMBP by 50% and C5 by 60% in the radiatum (**S.Fig. 10G-L**). Together, these findings indicate that WS exposure induces a distinct and attenuated neurobiological response characterized by altered amyloid balance and cell-state remodeling without overt oxidative or inflammatory injury.

## DISCUSSION

AirP exposure is increasingly recognized as a modifiable risk factor for AD, yet the biological mechanisms linking heterogeneous airborne pollutants to neurodegenerative pathology have remained unclear. Here, we demonstrate that acute exposure to two chemically distinct but metal-rich forms of AirP, DEP, and WTC dust elicit a highly conserved molecular response characterized by ferroptotic priming, impaired lipid peroxide detoxification, enhanced amyloidogenic processing, and selective white-matter vulnerability. Despite major differences in particle size, carbon content, and ionic composition, DEP and WTC converge on shared transcriptional, biochemical, and microstructural changes relevant to AD. In contrast, WS, produced by the incomplete combustion of wood with a substantially lower metal burden, induces a divergent and attenuated response that does not recapitulate these convergent features. Together, these findings identify metal-associated oxidative mechanisms as a unifying link between specific forms of AirP and AD-relevant pathology.

A central insight from this study is that chemical heterogeneity does not preclude biological convergence. DEP and WTC dust differ profoundly in origin and composition, with DEP representing PM_0.1_, carbon-rich traffic-related pollution, and WTC dust representing PM_10_, mineral-dominated particulate matter. Yet both exposures robustly activate overlapping transcriptomic responses. Approximately 1,200 genes were regulated in the same direction following DEP and WTC exposure, with strong rank–rank correlation across the transcriptome. These shared responses encompassed pathways related to oxidative stress, interferon signaling, DNA repair, and protein trafficking, suggesting engagement of a conserved stress-adaptive response to metal-rich AirP. The absence of comparable overlap with WS indicates that this response is not a generic consequence of particulate inhalation but rather suggests a shared biological response driven by metal content and redox activity.

Biochemically, DEP and WTC exposures produced a pattern consistent with ferroptotic priming rather than overt ferroptotic cell death. Both exposures increased lipid peroxidation while simultaneously impairing PCOOH detoxification, particularly through suppression of GPx4 and Prdx6-mediated activity. Importantly, these functional deficits occurred despite compensatory increases in antioxidant protein abundance, suggesting the increased protein is compensatory. Such decoupling is a defining feature of ferroptosis-vulnerable states and closely mirrors alterations reported in postmortem AD brain tissue with reduced GSH, PCOOH activity, and increased lipid peroxidation^26,27,32^. Concurrent reduction in FSP1 with WTC dust, and disruption of lipid remodeling pathways further support the interpretation that acute metal-rich AirP exposure sensitizes neural membranes to oxidative injury.

Lipid rafts are a critical nexus linking ferroptotic events to amyloidogenic processing. These membrane microdomains are enriched in polyunsaturated lipids and harbor the enzymatic machinery responsible for APP cleavage ^27,67,68^. Acute DEP and WTC exposure impaired lipid raft-specific antioxidant defenses, reduced phospholipid hydroperoxide detoxification capacity, and promoted accumulation of lipid peroxidation products. In parallel, both exposures increased amyloid-β production, with a preferential rise in the aggregation-prone Aβ42 species. These findings support previous cell culture experiments with TRAP^67^ and a model in which metal-driven lipid peroxidation within lipid rafts alters membrane composition and secretase activity^35,36^, thereby biasing APP processing toward amyloidogenic pathways. Notably, this constellation of lipid raft oxidative damage, impaired GPx4 activity, and increased Aβ42 production closely parallels biochemical signatures observed in human AD cortex^26,27^.

Iron handling and heme metabolism were selectively altered by DEP and WTC exposure, reinforcing the role of redistributed redox-active iron rather than bulk iron accumulation. Although total brain iron levels were unchanged acutely, both exposures disrupted ferritin storage capacity with increased tissue heme for DEP. Induction of HMOX1, particularly following WTC exposure, suggests enhanced heme turnover and liberation of redox-active iron, consistent with vascular stress and micro hemorrhagic processes reported in AirP-exposed and AD populations^24,69–72^. These findings align with neuropathological and imaging evidence linking cerebral microbleeds, iron redistribution, and AD progression^22,26,27,73–75^, indicating that even transient perturbations in iron handling may be sufficient to initiate lipid peroxidation cascades.

The structural consequences of these molecular changes were evident in selective white-matter vulnerability, most prominently within the corpus callosum. Both DEP and WTC exposure altered DTI MRI metrics, characterized by increased fractional anisotropy and reduced mean diffusivity. This pattern is consistent with acute microstructural alterations such as myelin-associated swelling, altered axonal packing, or changes in extracellular water diffusion rather than overt tissue loss^76,77^. DEP uniquely induced lipid and DNA oxidation within the corpus callosum, whereas WTC altered diffusion properties in the absence of overt oxidative damage, suggesting that microstructural disruption may precede detectable biochemical injury. These findings are consistent with emerging evidence that white matter is an early and sensitive target of environmental insults and may represent a substrate for later cognitive decline^78^.

In contrast to DEP and WTC, WS exposure produced a fundamentally different biological response. WS contained substantially lower metal content and did not induce coordinated ferroptotic priming, lipid peroxidation, or amyloidogenic bias. Although WS impaired phospholipid peroxide detoxification activity, this occurred in the absence of lipid peroxidation or ferroptosis-associated transcriptional responses. WS instead favored increased Aβ40 production without elevating Aβ42, consistent with a less pathogenic amyloid profile^79,80^. WS however has been reported to These findings underscore that not all particulate exposures carry equivalent neurodegenerative risk, and that metal burden is a key determinant of pathogenic potential.

Several limitations should be acknowledged. First, this study focused on acute exposures to capture early mechanistic responses rather than cumulative pathology. Behavioral assessments were intentionally not performed, as the acute exposure paradigm was designed to preserve post-exposure molecular, biochemical, and imaging endpoints that would be confounded by behavioral testing. While acute priming does not equate to disease, experimental evidence from independent exposure paradigms indicates that increasing duration or repetition of metal-rich air pollution exposure is associated with amplified lipid peroxidation, inflammatory activation, white-matter injury, and amyloid-related pathology. Together with the present findings, these data support a model in which repeated exposure to metal-rich AirP reinforces ferroptotic vulnerability, lipid raft oxidation, and amyloidogenic bias, thereby accelerating AD-associated pathology. Second, WS exposure was conducted at higher concentrations and in a single sex, reflecting real-world wildfire conditions but limiting direct quantitative comparison across exposures^43^.

In summary, this study identifies metal-driven oxidative vulnerability as a convergent biological mechanism linking chemically distinct forms of air pollution to AD-relevant pathology. Acute exposure to DEP and WTC dust primes the brain for ferroptotic stress, disrupts lipid raft antioxidant defenses, enhances amyloidogenic processing, and selectively alters white-matter microstructure, all of which are pathological features of AD. These effects are not shared by metal-poor WS, demonstrating that neurodegenerative risk is not an inevitable consequence of particulate exposure but depends on specific chemical features. By delineating a shared, mechanistically coherent pathway connecting AirP to AD, these findings provide a framework for understanding environmental contributions to neurodegeneration and highlight metal handling and lipid peroxide detoxification as actionable targets for intervention.

## Abbreviations

Aβ: amyloid-β
AhR: aryl hydrocarbon receptor
APP: amyloid precursor protein
ARNT: aryl hydrocarbon receptor nuclear translocator
BACE1: β-site amyloid precursor protein cleaving enzyme 1
DEP: diesel exhaust particles
FSP1: ferroptosis suppressor protein 1
GPx4: glutathione peroxidase 4
GSH: glutathione
HMOX1: heme oxygenase 1
NCOA4: nuclear receptor coactivator 4
WTC: World Trade Center dust

## Ethics Declarations

WJM reports the following: consultant: Viseon, Imperative Care, Q’Apel, Medtronic, Stryker, Stream Biomedical, Spartan Micro; Egret. Investor: Cerebrotech, Q’ Apel, Endostream, Viseon, Rebound, Stream Biomedical, Spartan Micro, Radical Catheters, Vastrax, Borvo

## Data Availability

The authors declare that the data supporting the findings of this study are available within the paper and its Supplementary Information files. Should any raw data files be needed in another format, they are available from the corresponding author upon reasonable request.

## Acknowledgements

Venn diagrams were made with Venny 2.1.0, and schematics were made using BioRender.

---

**Supplemental Figure 1:** Chemical composition of DEP and WTC dust showing differences in the chemical composition aside from the metal content. **A)** Water-soluble inorganic ions quantified by ion chromatography. **B)** Carbonaceous fraction (elemental carbon, EC; organic carbon, OC) measured by thermal optical analysis. Metal content measured by ICP-MS after acid digestion of particulate samples for **C)** DEP and WTC, and **D)** WS (n=18).

**Supplemental Figure 2:** Volcano plots showing sex-stratified DEGs for cortex exposed acutely to DEP or WTC dust compared to FA. **A)** DEP vs FA Male, **B)** DEP vs FA Female, **C)** DEP Female vs DEP Male, **D)** WTC vs FA Male, **E)** WTC vs FA Female, **F)** WTC Female vs WTC Male, **G)** DEP vs WTC Male, and **H)** DEP vs WTC Female.

**Supplemental Figure 3:** Woodsmoke has limited overlap with DEP and WTC. **A)** Volcano plots for WS gene responses from FA. **B)** GSEA pathway enrichment for WS exposure. Rank-Rank plots DEP vs WS by **C)** gene and **D)** hex plot for gene density. Rank-Rank plots WTC vs WS by **E)** gene and **F)** hex plot for gene density. **G)** Upset plot showing the number of DEGs exclusive or shared between the DEP, WTC, and WS exposures.

**Supplemental Figure 4:** Responses to copper handling and acute woodsmoke exposure do not impact ferroptotic-related antioxidants. Cortical levels of **A)** HNE by IF (all cortical layers stitched at 40x) and **B)** NT by dot blot for DEP and WTC. WS analyses of antioxidants by Western and dot blot for **C)** HNE, **D)** NT, **E)** GPx4, **F)** Prdx6, **G)** PCOOH activity, **H)** GPx1, **I)** total GPx activity, **J)** FSP1, **K)** xCT, **L)** GCLM, **M)** GCLC, N) GSTA4, **O)** ALDH2, and **P)** ACSL4 measured in whole cell lysates. **Q)** Copper signaling pathways, **R)** comparison of copper-related genes for DEP, WTC, and WS. Western blots for **S)** CCS and **T)** SOD1 and **U)** SOD activity for DEP and WTC. Western blot of **V)** SOD1 and **W)** SOD activity for WS. Statistical analysis was performed using ANCOVA adjusted for sex with Bonferroni’s post hoc test for A,B,S-U, or two-tailed t-test for C-P,V,W. *p<0.05, **p<0.01, ***p<0.001.

**Supplemental Figure 5:** DNA damage but not RNA errors increase with DEP. **A)** Heatmap showing DEGs involved in DNA damage response, base excision repair, and RNA surveillance pathways in the cortex following acute exposure to DEP or WTC dust. **B)** Immunofluorescence quantification of cortical 8-OHdG (all cortical layers stitched at 40x). Error rate and spectrum of transcript errors for **C)** RNApol I, **D)** RNApol II, **E)** RNApol III, and **D)** RNApol Mito from cortical RNA for FA, DEP, and WTC males (n=1). Statistical analysis was performed using ANCOVA adjusted for sex with Bonferroni’s post hoc test. *p<0.05.

**Supplemental Figure 6:** Brain iron levels are not altered by AirP. *Ex vivo* multi-echo multi-slice MRI by **A)** brain region and by **B)** white and grey matter in acutely exposed DEP and WTC exposed brains. Western blots and assays for iron signaling responses to WS exposure: **C)** TF, **D)** TfR, **E)** DMT1, **F)** FTL, **G)** FTH1, **H)** FPN, **I)** tissue heme, **J)** HCP1, **K)** HMOX1, **L)** IRP1, and **M)** IRP2. Statistical analysis was performed using two-way ANOVA for A,B, or two-tailed t-test for C-M. **p<0.01, ***p<0.001.

**Supplemental Figure 7:** WS exposure increases soluble Aβ40 but not Aβ42. Western or dot blot for **A)** APP, **B)** ADAM10, **C)** BACE1, **D)** PSEN1, **E)** Aβ40, **F)** Aβ42, **G)** ApoE, and **H)** LRP1 in cortex from RIPA lysates. Statistical analysis was performed using two-tailed t-test, *p<0.05, **p<0.01, ***p<0.00

**Supplemental Figure 8:** Transcription factor responses to acute woodsmoke exposure. Western blots of nuclear lysates for **A)** AhR, **B)** ARNT1, **C)** ARNT2, **D)** NCOA4, **E)** BACH1, **F)** Nrf2, **G)** ATF4, and **H)** NF-κB p65. Statistical analysis was performed using two-tailed t-test, *p<0.05, **p<0.01.

**Supplemental Figure 9:** Cortical oligodendrocyte and neuron proportions are shifted with WS exposure, opposite of DEP and WTC changes. Western blots for **A)** MBP and **B)** NeuN in DEP and WTC-exposed mice. Cell type proportions for **C)** oligodendrocytes and **D)** neurons. Western blots for **E)** MBP, **F)** NeuN, DoRothEA transcription factor activity, and Western blot of nuclear **G)** NeuroD1. Statistical analysis was performed using ANCOVA adjusted for sex with Bonferroni’s post hoc test for A,B. ANCOVA with Kruskal-Wallis for C,D, and two-tailed t-test for E,F, and H. **p<0.01.

**Supplemental Figure 10:** Oxidative damage and inflammatory markers are not altered by acute woodsmoke exposure in the corpus callosum or hippocampus. Immunofluorescence analysis of the corpus callosum showing **A)** HNE, **B)** 8-OHdG, **C)** dMBP, **D)** C5, **E)** C5a, and **F)** Iba1. Representative images are of 40x corpus callosum region (the same animal was selected as the representative image for: HNE/Iba-1 in FA/WS; C5/dMBP in FA; 8-OHdG/C5/dMBP in WS. Multiple stains were performed on the same slices). Immunofluorescence analysis of hippocampal subregions, including the granule cell layer, molecular layer, and polymorphic layer of the dentate gyrus, as well as the stratum oriens and stratum radiatum of the cornu ammonis, showing **G)** HNE, **H)** 8-OHdG, **I)** dMBP, **J)** C5, **K)** C5a, and **L)** Iba1. Statistical analysis was performed using two-tailed t-tests, **p<0.01, ****p<0.0001.

## References

1. Estimation of the global prevalence of dementia in 2019 and forecasted prevalence in 2050: an analysis for the Global Burden of Disease Study 2019 - The Lancet Public Health. https://www.thelancet.com/journals/lanpub/article/PIIS2468-2667(21)00249-8/fulltext.

2. Karran, E., Mercken, M. & Strooper, B. D. The amyloid cascade hypothesis for Alzheimer’s disease: an appraisal for the development of therapeutics. Nat Rev Drug Discov 10, 698–712 (2011).

3. Glabe, C. G. Structural Classification of Toxic Amyloid Oligomers*. Journal of Biological Chemistry 283, 29639–29643 (2008).

4. Haghani, A., Thorwald, M., Morgan, T. E. & Finch, C. E. The APOE gene cluster responds to air pollution factors in mice with coordinated expression of genes that differs by age in humans. Alzheimers Dement Feb;17(2):175–190, (2021).

5. Franz, C. E., Gustavson, D. E., Elman, J. A. & Fennema-Notestine, C. Associations Between Ambient Air Pollution and Cognitive Abilities from Midlife to Early Old Age: Modification by APOE Genotype. J Alzheimers Dis 93, 193–209 (2023).

6. Finch, C. E. & Kulminski, A. M. The Alzheimer’s Disease Exposome. Alzheimers Dement Sep;15(9):1123**-**1132, (2019).

7. Finch, C. & Thorwald, M. Inhaled pollutants of the Gero-Exposome and later life health. The journals of gerontology. Series A, Biological sciences and medical sciences 10.1093/gerona/glae107 (2024) doi:10.1093/gerona/glae107.

8. Cacciottolo, M. et al. Particulate air pollutants, APOE alleles and their contributions to cognitive impairment in older women and to amyloidogenesis in experimental models. Translational Psychiatry 7, e1022–e1022 (2017).

9. Wang, X. et al. Ambient Air Pollution and Long-Term Trajectories of Episodic Memory Decline among Older Women in the WHIMS-ECHO Cohort. Environ Health Perspect 129, 97009 (2021).

10. Kilian, J. & Kitazawa, M. The emerging risk of exposure to air pollution on cognitive decline and Alzheimer’s disease – Evidence from epidemiological and animal studies. Biomed J 41, 141–162 (2018).

11. Calderón-Garcidueñas, L. et al. Interactive and additive influences of Gender, BMI and Apolipoprotein 4 on cognition in children chronically exposed to high concentrations of PM2.5 and ozone. APOE 4 females are at highest risk in Mexico City. Environ Res 150, 411–422 (2016).

12. Zhang, H. et al. Urban Air Pollution Nanoparticles from Los Angeles: Recently Decreased Neurotoxicity. J Alzheimers Dis 82, 307–316 (2021).

13. Cory-Slechta, D. A., Merrill, A. & Sobolewski, M. Air Pollution-Related Neurotoxicity Across the Life Span. Annu Rev Pharmacol Toxicol 63, 143–163 (2023).

14. Iban-Arias, R. et al. Exposure to World Trade Center Dust Exacerbates Cognitive Impairment and Evokes a Central and Peripheral Pro-Inflammatory Transcriptional Profile in an Animal Model of Alzheimer’s Disease. Journal of Alzheimer’s Disease 91, 779–794 (2023).

15. Xue, Y. et al. Environmental aluminum oxide inducing neurodegeneration in human neurovascular unit with immunity. Sci Rep 14, 744 (2024).

16. Calderón-Garcidueñas, L. et al. The impact of environmental metals in young urbanites’ brains. Exp Toxicol Pathol 65, 503–511 (2013).

17. Smith, M. A., Harris, P. L. R., Sayre, L. M. & Perry, G. Iron accumulation in Alzheimer disease is a source of redox-generated free radicals. Proceedings of the National Academy of Sciences 94, 9866–9868 (1997).

18. Ashraf, A., Jeandriens, J., Parkes, H. G. & So, P.-W. Iron dyshomeostasis, lipid peroxidation and perturbed expression of cystine/glutamate antiporter in Alzheimer’s disease: Evidence of ferroptosis. Redox Biol 32, 101494 (2020).

19. Dedman, D. J. et al. Iron and aluminium in relation to brain ferritin in normal individuals and Alzheimer’s-disease and chronic renal-dialysis patients. Biochem J 287, 509–514 (1992).

20. Adivi, A., Lucero, J., Simpson, N., McDonald, J. D. & Lund, A. K. Exposure to Traffic-Generated Air Pollution Promotes Alterations in the Integrity of the Brain Microvasculature and Inflammation in Female ApoE−/− mice. Toxicol Lett 339, 39–50 (2021).

21. Hartz, A. M. S., Bauer, B., Block, M. L., Hong, J.-S. & Miller, D. S. Diesel exhaust particles induce oxidative stress, proinflammatory signaling, and P-glycoprotein up-regulation at the blood-brain barrier. FASEB J 22, 2723–2733 (2008).

22. Nag, S. et al. Ex vivo MRI facilitates localization of cerebral microbleeds of different ages during neuropathology assessment. Free Neuropathology 2, 35–35 (2021).

23. Yates, P. A. et al. Incidence of cerebral microbleeds in preclinical Alzheimer disease. Neurology 82, 1266–1273 (2014).

24. Shkirkova, K., Demetriou, A. N. & Sizdahkhani, S. Microglial TLR4 Mediates White Matter Injury in a Combined Model of Diesel Exhaust Exposure and Cerebral Hypoperfusion. Stroke 55, 1090–1093 (2024).

25. Halliwell, B., Adhikary, A., Dingfelder, M. & Dizdaroglu, M. Hydroxyl radical is a significant player in oxidative DNA damage in vivo. Chemical Society reviews 50, 8355 (2021).

26. Thorwald, M. A. et al. Iron-associated lipid peroxidation in Alzheimer’s disease is increased in lipid rafts with decreased ferroptosis suppressors, tested by chelation in mice. Alzheimer’s & Dementia 21, e14541 (2025).

27. Thorwald, M. A., et al. Down syndrome with Alzheimer’s disease brains have increased iron and associated lipid peroxidation consistent with ferroptosis. Alzheimer’s & Dementia 21, e70322 (2025).

28. Buxton, G. V. & Elliot, A. J. Rate constant for reaction of hydroxyl radicals with bicarbonate ions. International Journal of Radiation Applications and Instrumentation. Part C. Radiation Physics and Chemistry 27, 241–243 (1986).

29. Dixon, S. J. et al. Ferroptosis: an iron-dependent form of nonapoptotic cell death. Cell 149, 1060–1072 (2012).

30. Ayton, S. et al. Regional brain iron associated with deterioration in Alzheimer’s disease: A large cohort study and theoretical significance. Alzheimer’s & Dementia 17, 1244–1256 (2021).

31. Lane, D. J. R., Alves, F., Ayton, S. & Bush, A. I. Striking a NRF2: The rusty and rancid vulnerabilities toward ferroptosis in Alzheimer’s disease. Antioxid Redox Signal 10.1089/ars.2023.0318 (2023) doi:10.1089/ars.2023.0318.

32. Mandal, P. K., Saharan, S., Tripathi, M. & Murari, G. Brain glutathione levels--a novel biomarker for mild cognitive impairment and Alzheimer’s disease. Biol Psychiatry 78, 702–710 (2015).

33. Zhang, H., Morgan, T. E. & Forman, H. J. Age-related alteration in HNE elimination enzymes. Arch Biochem Biophys 699, 108749 (2021).

34. Castro, J. P., Jung, T., Grune, T. & Siems, W. 4-Hydroxynonenal (HNE) modified proteins in metabolic diseases. Free Radical Biology and Medicine 111, 309–315 (2017).

35. Kwak, Y.-D. et al. Differential regulation of BACE1 expression by oxidative and nitrosative signals. Molecular Neurodegeneration 6, 17 (2011).

36. Chen, L., Na, R., Gu, M., Richardson, A. & Ran, Q. Lipid peroxidation upregulates BACE1 expression in vivo: a possible early event of amyloidogenesis in Alzheimer’s disease. J Neurochem 107, 197–207 (2008).

37. Godoy-Lugo, J. A. et al. Air pollution decreases postsynaptic PSD-95 and NMDA receptor subunits in synaptosomes from mouse cerebral cortex. Environ Pollut 126845 (2025) doi:10.1016/j.envpol.2025.126845.

38. Godoy-Lugo, J. A. et al. Air pollution amyloidogenesis is attenuated by the gamma-secretase modulator GSM-15606. Alzheimers Dement 10.1002/alz.14086 (2024) doi:10.1002/alz.14086.

39. Clouston, S. A. P. et al. Incidence of Dementia Before Age 65 Years Among World Trade Center Attack Responders. JAMA Network Open 7, e2416504 (2024).

40. Huang, C. et al. World Trade Center Site Exposure Duration Is Associated with Hippocampal and Cerebral White Matter Neuroinflammation. Mol Neurobiol 60, 160–170 (2023).

41. McGee, J. K. et al. Chemical analysis of World Trade Center fine particulate matter for use in toxicologic assessment. Environ Health Perspect 111, 972–980 (2003).

42. Lippmann, M., Cohen, M. D. & Chen, L.-C. Health effects of World Trade Center (WTC) Dust: An unprecedented disaster’s inadequate risk management. Crit Rev Toxicol 45, 492–530 (2015).

43. Scieszka, D. et al. Neurometabolomic impacts of wood smoke and protective benefits of anti-aging therapeutics in aged female C57BL/6J mice. Part Fibre Toxicol 22, 23 (2025).

44. Scieszka, D. et al. Biomass smoke inhalation promotes neuroinflammatory and metabolomic temporal changes in the hippocampus of female mice. J Neuroinflammation 20, 192 (2023).

45. Cunningham, C. X. et al. Climate-linked escalation of societally disastrous wildfires. Science 390, 53–58 (2025).

46. Chow, J. C. et al. The IMPROVE_A temperature protocol for thermal/optical carbon analysis: Maintaining consistency with a long-term database. Journal of the Air and Waste Management Association 57, 1014–1023 (2007).

47. Karthikeyan, S. & Balasubramanian, R. Determination of water-soluble inorganic and organic species in atmospheric fine particulate matter. Microchemical Journal 82, 49–55 (2006).

48. Watson, J. G. Ion chromatography in elemental analysis of airborne particles. https://www.researchgate.net/publication/235341492 (1999).

49. Herner, J. D., Green, P. G. & Kleeman, M. J. Measuring the trace elemental composition of size-resolved airborne particles. Environmental Science and Technology 40, 1925–1933 (2006).

50. Lough, G. C. et al. Emissions of metals associated with motor vehicle roadways. Environmental Science and Technology 39, 826–836 (2005).

51. Alander, T. J. A., Leskinen, A. P., Raunemaa, T. M. & Rantanen, L. Characterization of diesel particles: effects of fuel reformulation, exhaust aftertreatment, and engine operation on particle carbon composition and volatility. Environ Sci Technol 38, 2707–2714 (2004).

52. Barnes, S. R. et al. ROCKETSHIP: a flexible and modular software tool for the planning, processing and analysis of dynamic MRI studies. BMC Medical Imaging 15, 19 (2015).

53. Yeh, F.-C. DSI Studio: an integrated tractography platform and fiber data hub for accelerating brain research. Nat Methods 22, 1617–1619 (2025).

54. Thorwald, M. A., Silva, J., Head, E. & Finch, C. E. Amyloid futures in the expanding pathology of brain aging and dementia. Alzheimers Dement 10.1002/alz.12896 (2022) doi:10.1002/alz.12896.

55. Godoy-Lugo, J. A. et al. Increase of brain Aβ peptides and secretase activity during normal aging in rodent and human. GeroScience 10.1007/s11357-025-01926-w (2025) doi:10.1007/s11357-025-01926-w.

56. Stolwijk, J. M., Falls-Hubert, K. C., Searby, C. C., Wagner, B. A. & Buettner, G. R. Simultaneous detection of the enzyme activities of GPx1 and GPx4 guide optimization of selenium in cell biological experiments. Redox Biol 32, 101518 (2020).

57. Arevalo, J. A. et al. Age-related declines in mitochondrial Prdx6 contribute to dysregulated muscle bioenergetics. Redox Biol 86, 103808 (2025).

58. Dobin, A. et al. STAR: ultrafast universal RNA-seq aligner. Bioinformatics 29, 15–21 (2013).

59. Liao, Y., Smyth, G. K. & Shi, W. The R package Rsubread is easier, faster, cheaper and better for alignment and quantification of RNA sequencing reads. Nucleic Acids Res 47, e47 (2019).

60. Love, M. I., Huber, W. & Anders, S. Moderated estimation of fold change and dispersion for RNA-seq data with DESeq2. Genome Biol 15, 550 (2014).

61. Chung, C. S. et al. Transcript errors generate amyloid-like proteins in human cells. Nat Commun 15, 8676 (2024).

62. Symons, J. et al. The mutational landscape of SARS-CoV-2 provides new insight into viral evolution and fitness. Nat Commun 16, 6425 (2025).

63. Wang, X., Park, J., Susztak, K., Zhang, N. R. & Li, M. Bulk tissue cell type deconvolution with multi-subject single-cell expression reference. Nat Commun 10, 380 (2019).

64. Schraufnagel, D. E. The health effects of ultrafine particles. Exp Mol Med 52, 311–317 (2020).

65. McGee, J. K. et al. Chemical analysis of World Trade Center fine particulate matter for use in toxicologic assessment. Environ Health Perspect 111, 972–980 (2003).

66. Gwon, A.-R. et al. Oxidative lipid modification of nicastrin enhances amyloidogenic γ-secretase activity in Alzheimer’s disease. Aging Cell 11, 559–568 (2012).

67. Cacciottolo, M. et al. Traffic-related air pollutants (TRAP-PM) promote neuronal amyloidogenesis through oxidative damage to lipid rafts. Free Radical Biology and Medicine 147, 242–251 (2020).

68. Sezgin, E., Levental, I., Mayor, S. & Eggeling, C. The mystery of membrane organization: composition, regulation and roles of lipid rafts. Nat. Rev. Mol. Cell Biol. 18, 361–374 (2017).

69. Cotter, D. L. et al. Exposure to multiple ambient air pollutants changes white matter microstructure during early adolescence with sex-specific differences. Commun Med (Lond) 4, 155 (2024).

70. Kusters, M. S. W. et al. Residential ambient air pollution exposure and the development of white matter microstructure throughout adolescence. Environmental Research 262, 119828 (2024).

71. Akoudad, S. et al. Association of Cerebral Microbleeds With Cognitive Decline and Dementia. JAMA Neurol 73, 934–943 (2016).

72. Haller, S. et al. Radiologic-Histopathologic Correlation of Cerebral Microbleeds Using Pre-Mortem and Post-Mortem MRI. PLOS ONE 11, e0167743 (2016).

73. Damulina, A. et al. Cross-sectional and Longitudinal Assessment of Brain Iron Level in Alzheimer Disease Using 3-T MRI. Radiology 296, 619–626 (2020).

74. Ayton, S., Faux, N. G., Bush, A. I., & Alzheimer’s Disease Neuroimaging Initiative. Ferritin levels in the cerebrospinal fluid predict Alzheimer’s disease outcomes and are regulated by APOE. Nat Commun 6, 6760 (2015).

75. Ayton, S. et al. Brain iron is associated with accelerated cognitive decline in people with Alzheimer pathology. Mol Psychiatry 25, 2932–2941 (2020).

76. Lin, M., He, H., Schifitto, G. & Zhong, J. Simulation of Changes in Diffusion Related to Different Pathologies at Cellular Level After Traumatic Brain Injury. Magn Reson Med 76, 290–300 (2016).

77. Lee, J. K. et al. Fractional anisotropy from diffusion tensor imaging correlates with acute astrocyte and myelin swelling in neonatal swine models of excitotoxic and hypoxic-ischemic brain injury. J Comp Neurol 529, 2750–2770 (2021).

78. Garnier-Crussard, A., Cotton, F., Krolak-Salmon, P. & Chételat, G. White matter hyperintensities in Alzheimer’s disease: Beyond vascular contribution. Alzheimer’s & Dementia 19, 3738–3748 (2023).

79. Lambert, M. P. et al. Diffusible, nonfibrillar ligands derived from Aβ1–42 are potent central nervous system neurotoxins. Proceedings of the National Academy of Sciences 95, 6448–6453 (1998).

80. Jarrett, J. T., Berger, E. P. & Lansbury, P. T. The carboxy terminus of the beta amyloid protein is critical for the seeding of amyloid formation: implications for the pathogenesis of Alzheimer’s disease. Biochemistry 32, 4693–4697 (1993).

