## Supplementary figures and images for "Induction of ferroptotic and amyloidogenic signatures linked to Alzheimer’s disease by chemically distinct air pollutants"

### S.Fig. 1

#### Diesel Exhaust Particles & World Trade Center Dust

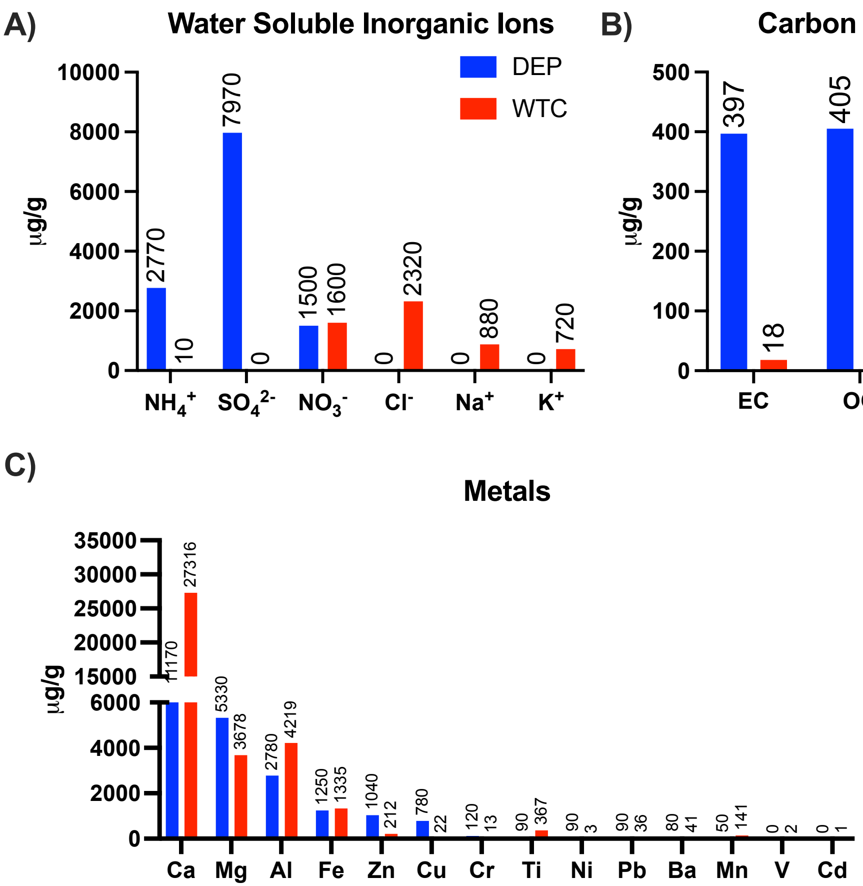


#### Woodsmoke

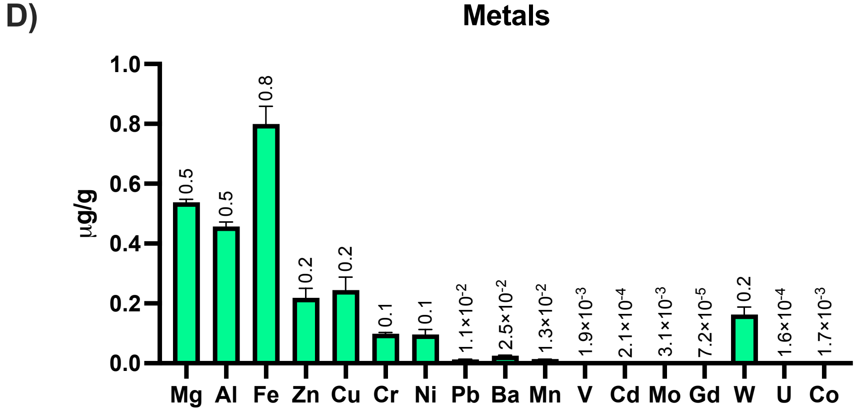

### S.Fig. 3

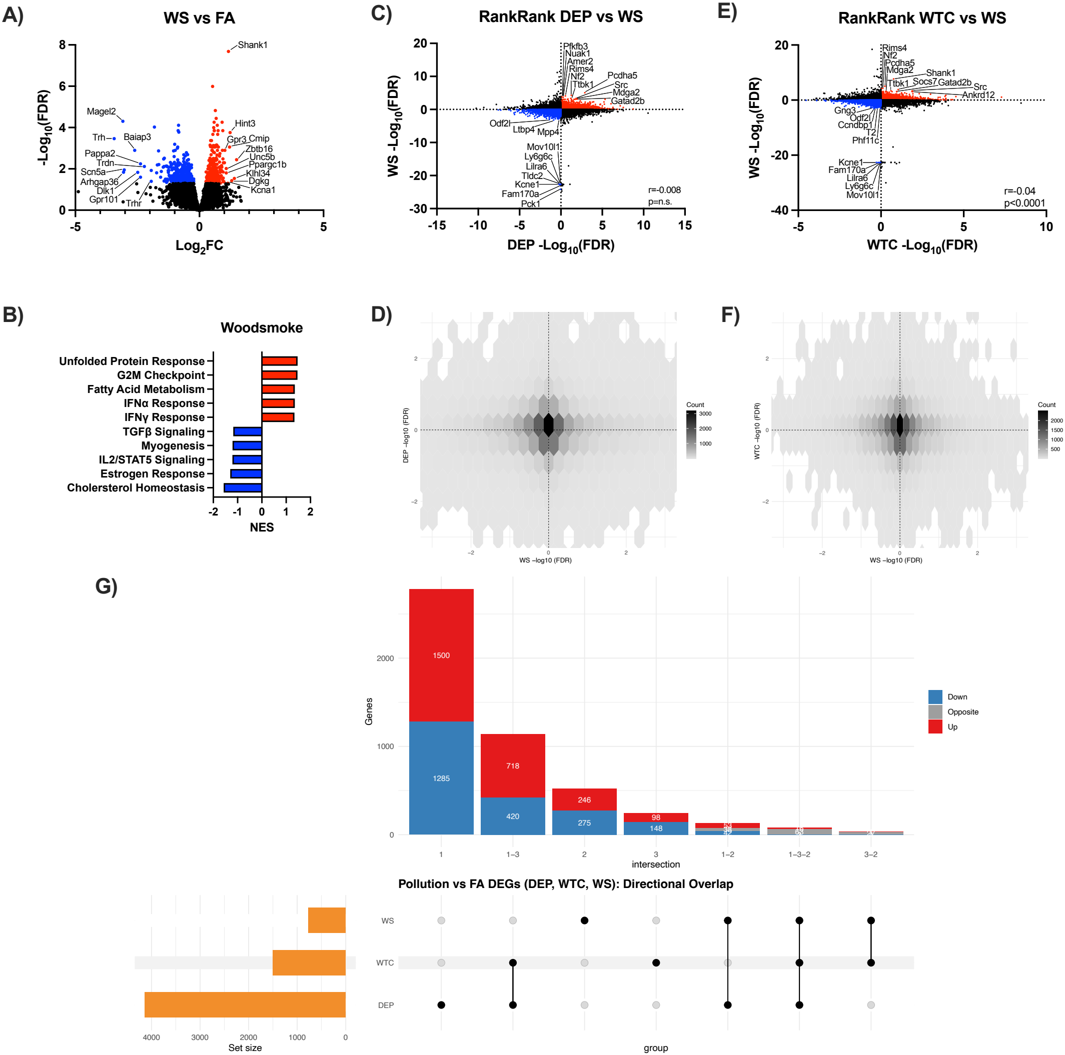

### S.Fig. 4

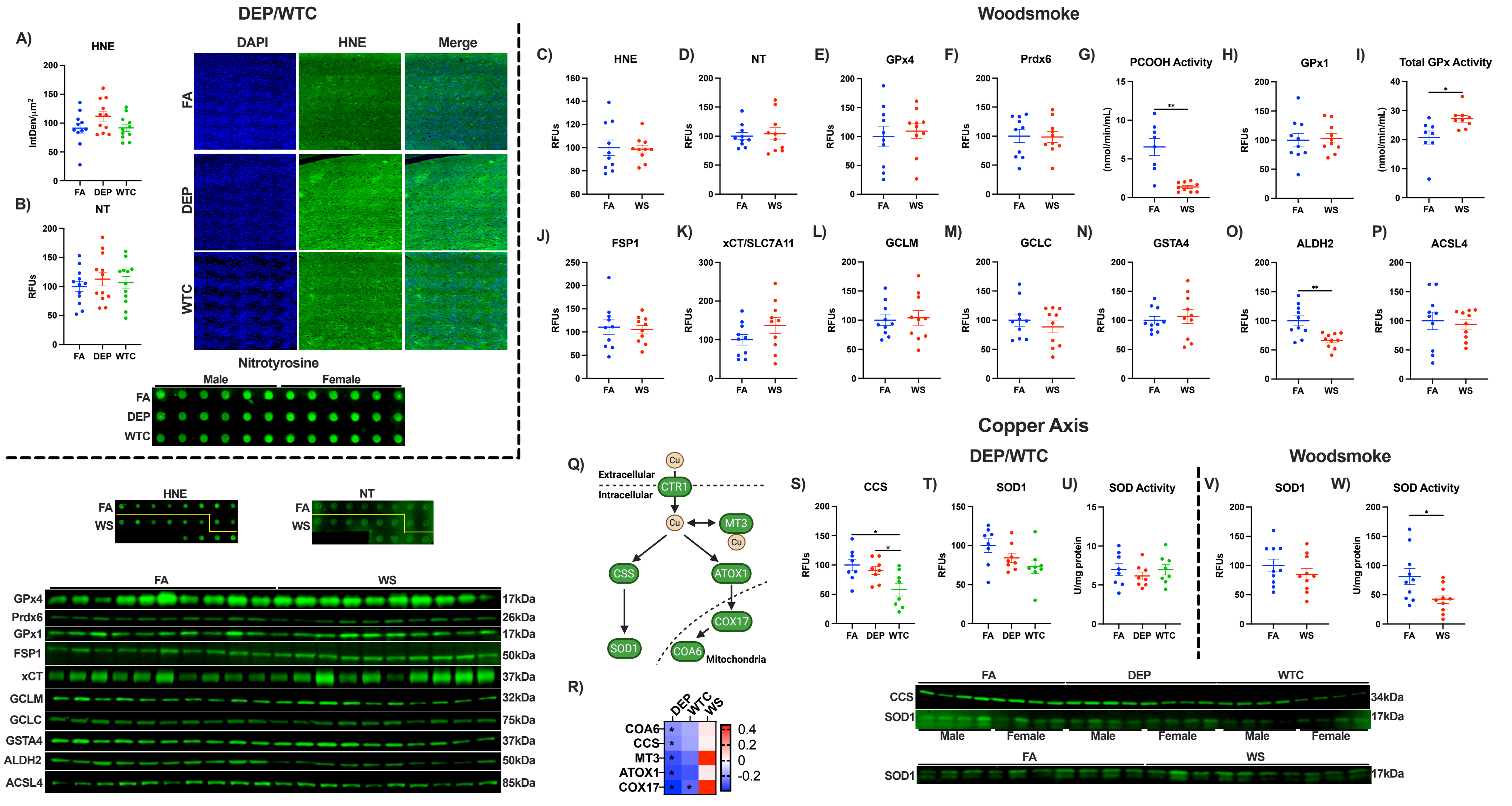

### S.Fig. 5

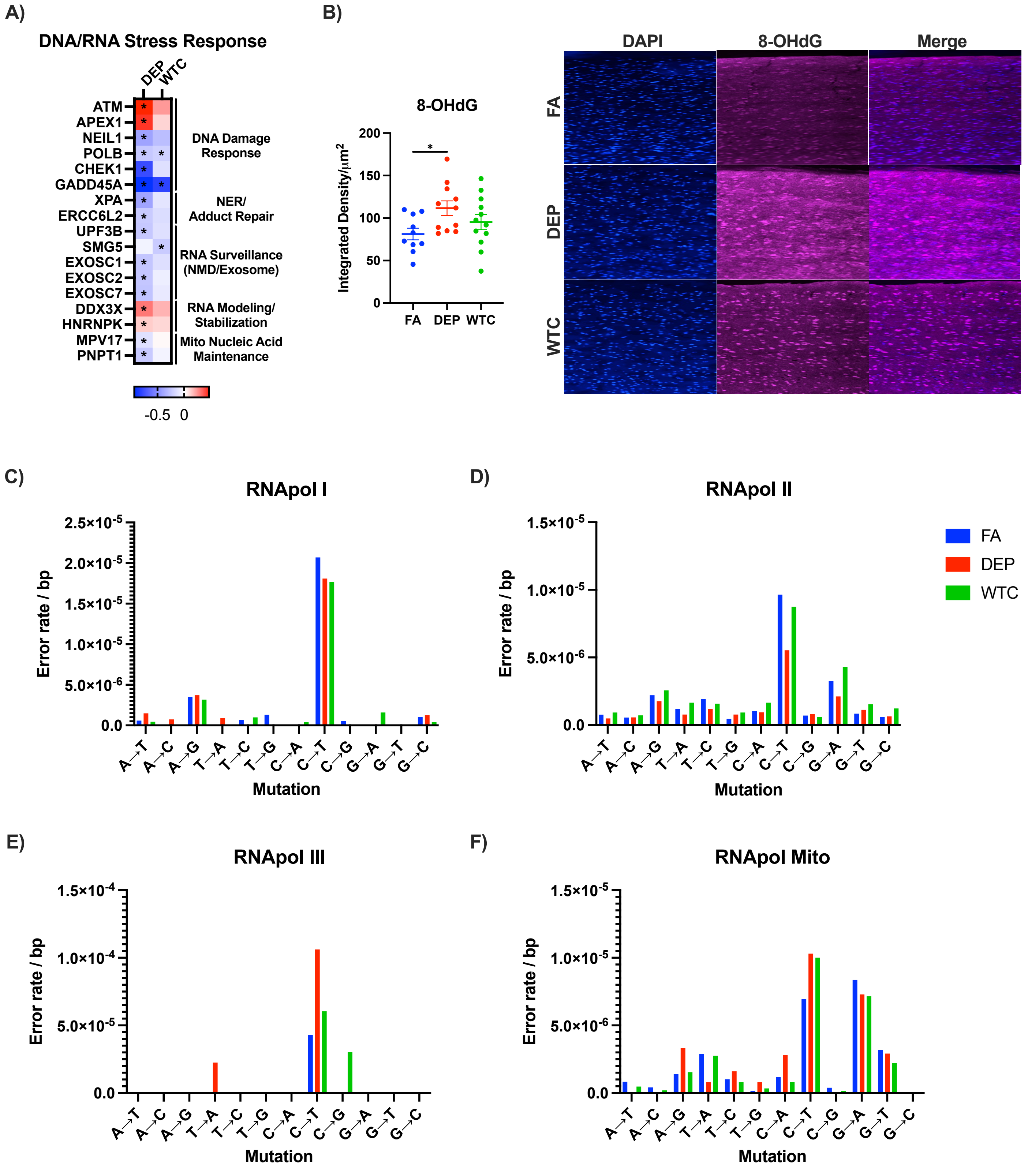

### S.Fig. 6

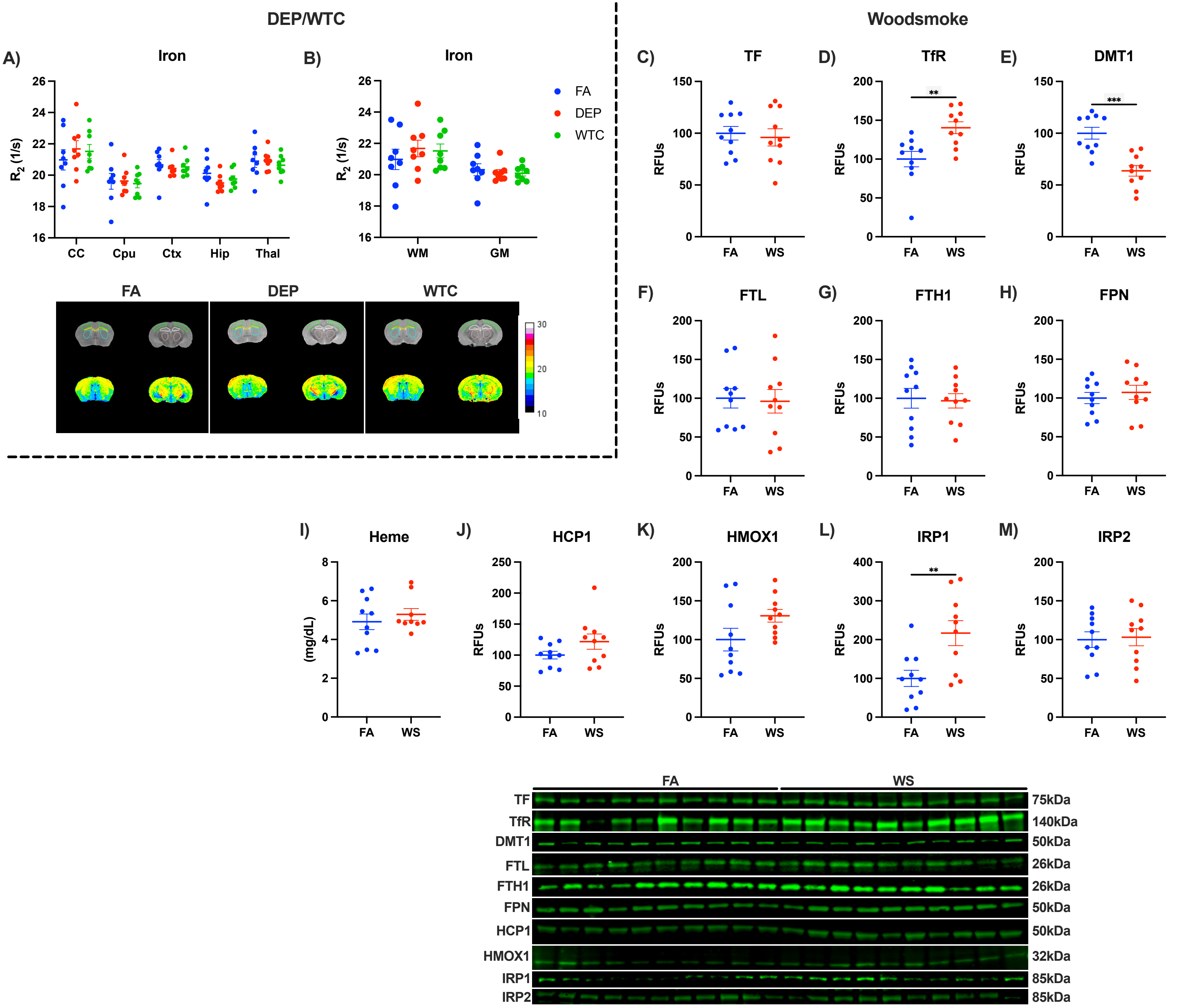

### S.Fig. 7

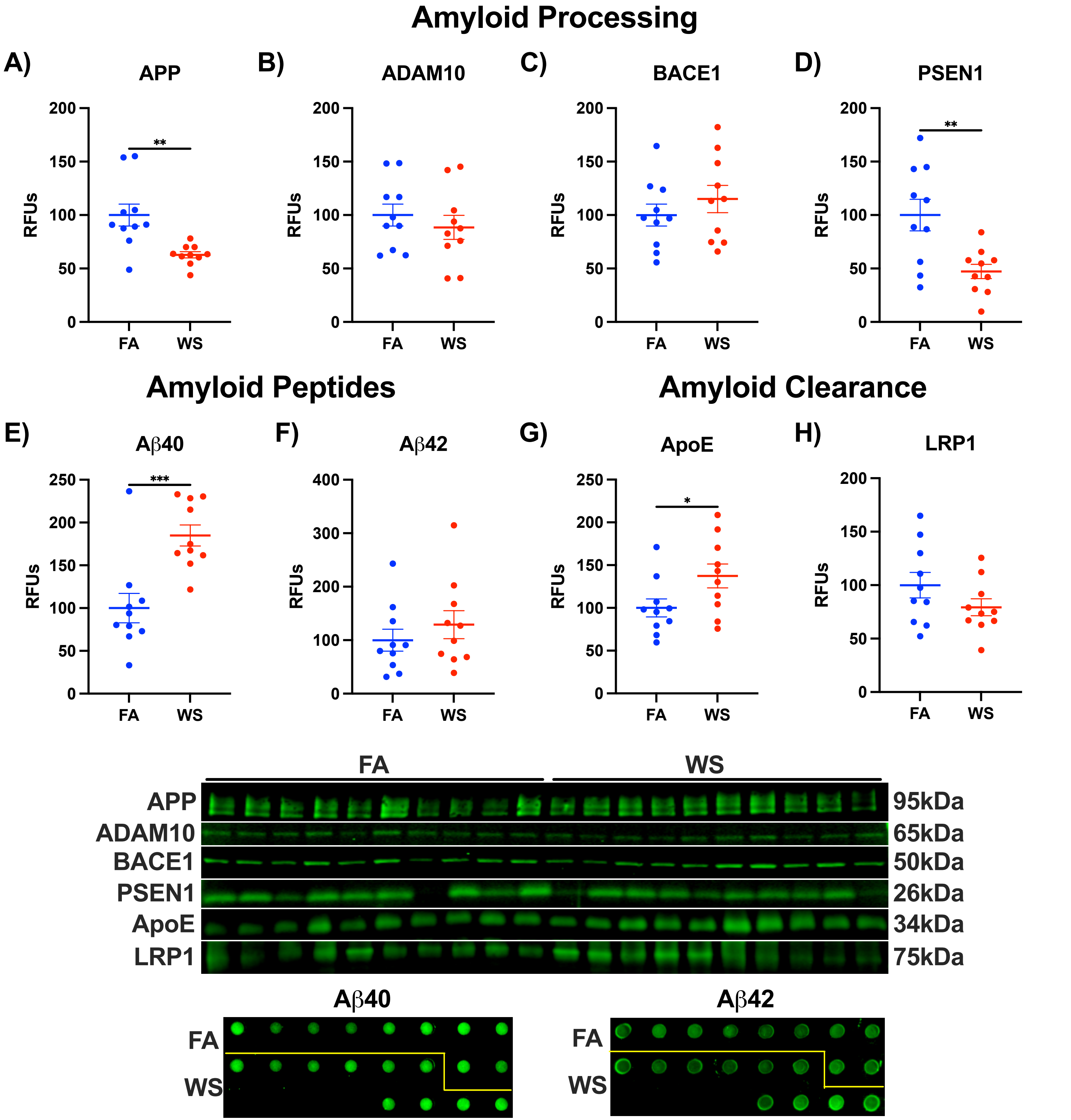

### S.Fig. 8

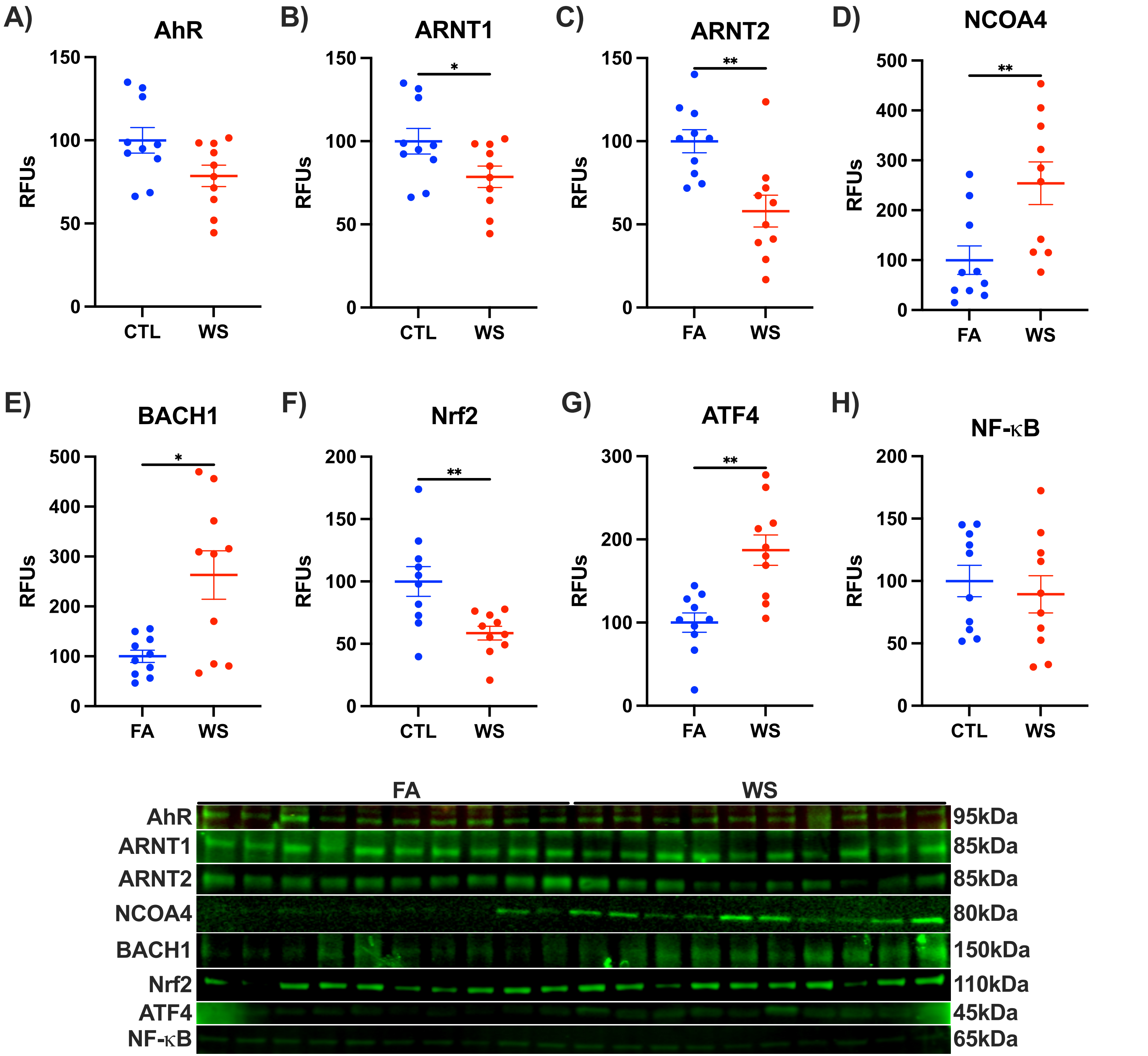

### S.Fig. 9

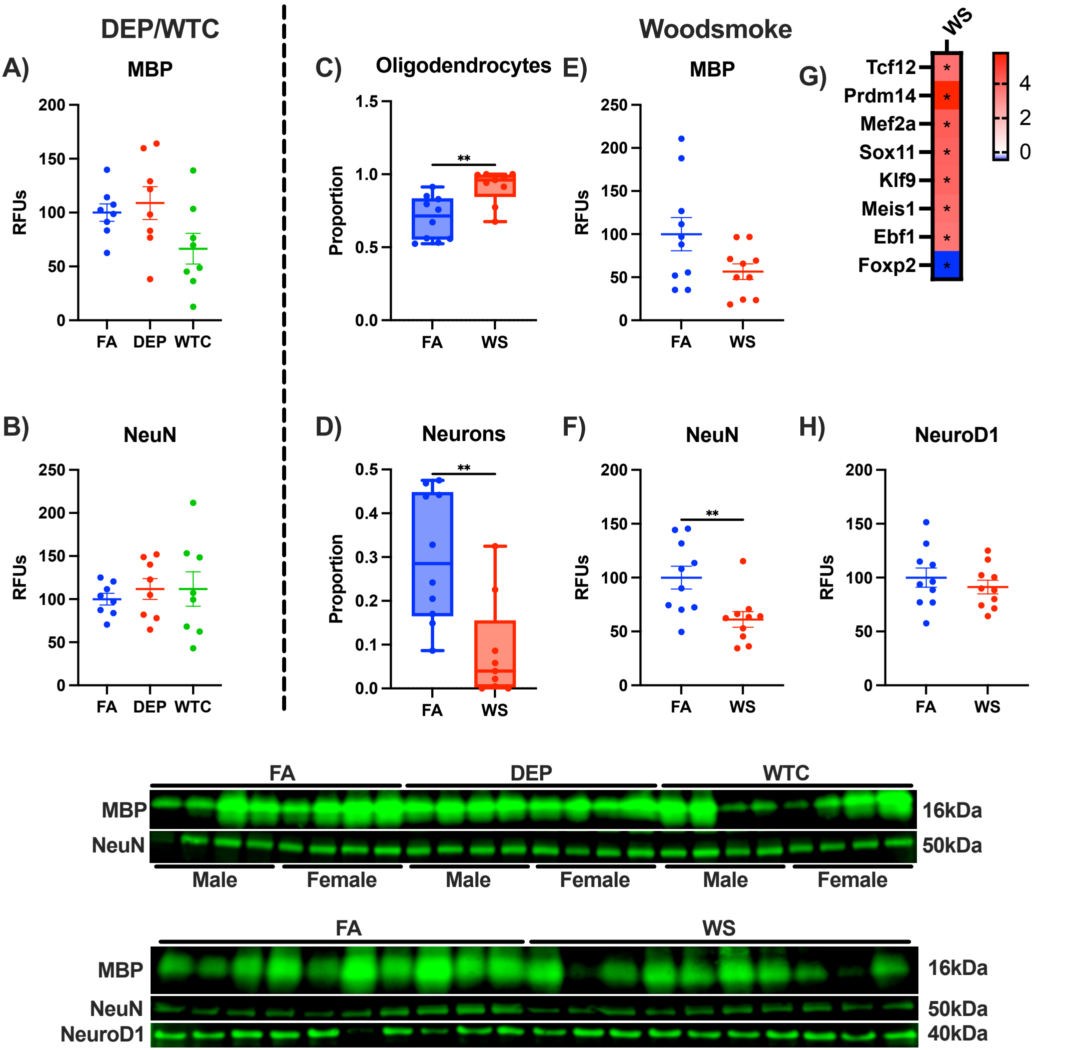

### S.Fig. 10

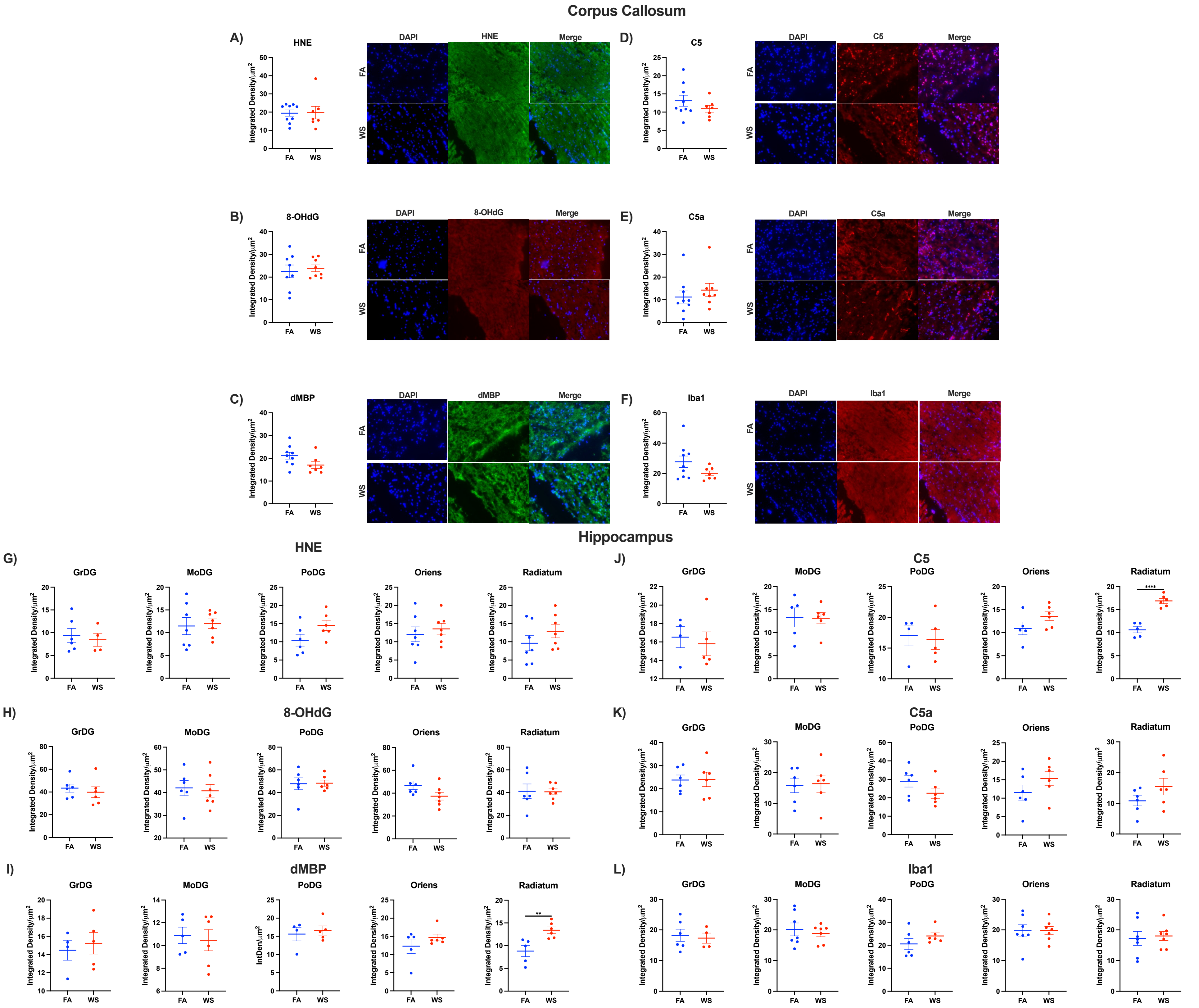

### Supplemental Figure 2

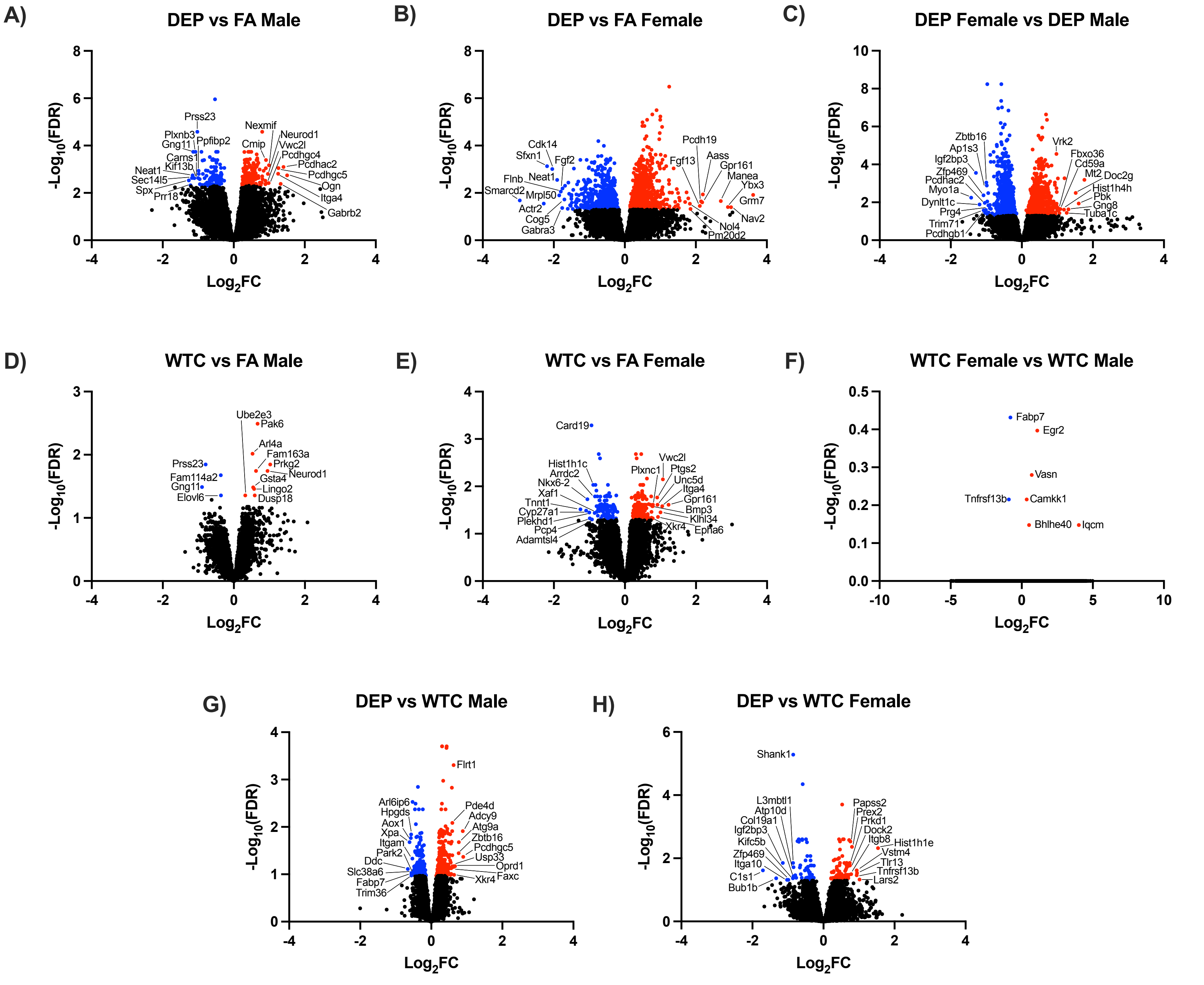
